# The AdcACB/AdcAII system is Essential for Zinc Homeostasis and an Important Contributor of *Enterococcus faecalis* Virulence

**DOI:** 10.1101/2021.12.01.470831

**Authors:** Ling Ning Lam, Debra N. Brunson, Jonathan J. Molina, Ana L. Flores-Mireles, José A. Lemos

**Author notes:** Address correspondence to José A. Lemos.

## Abstract

Bacterial pathogens require a variety of micronutrients for growth, including trace metals such as iron, manganese, and zinc (Zn). Despite their relative abundance in host environments, access to these metals is severely restricted during infection due to host-mediated defense mechanisms collectively known as nutritional immunity. Despite a growing appreciation of the importance of Zn in host-pathogen interactions, the mechanisms of Zn homeostasis and the significance of Zn to the pathophysiology of *E. faecalis*, a major pathogen of nosocomial and community-associated infections, have not been investigated. Here, we show that *E. faecalis* encoded an ABC-type transporter AdcACB and an orphan substrate-binding lipoprotein AdcAII that work cooperatively to maintain Zn homeostasis. Simultaneous inactivation of *adcA* and *adcAII* or the entire *adcACB* operon led to significant reduction in intracellular Zn under Zn-restricted conditions, heightened sensitivity to Zn-chelating agents including human calprotectin, aberrant cell morphology, and impaired fitness in serum *ex vivo*. Additionally, inactivation of *adcACB* and *adcAII* significantly reduced bacterial tolerance towards cell envelope-targeting antibiotics, which may be associated to altered fatty acid abundance and species. Lastly, we show that the AdcACB/AdcAII system contributes to *E. faecalis* virulence in an invertebrate (*Galleria mellonella*) infection model and in two catheter-associated mouse infection models that recapitulate many of the host conditions associated with enterococcal human infections. Collectively, this report reveals that high-affinity Zn import is essential for the pathogenesis of *E. faecalis* indicating that the surface-associated AdcA and AdcAII lipoproteins are potential therapeutic targets.

## INTRODUCTION

The first-row *d*-block elements iron (Fe), manganese (Mn), and zinc (Zn) are essential trace metals to all forms of life by serving structural, catalytic and regulatory functions to metalloproteins involved in a variety of biological processes (1–4). As a result of this essentiality, hosts deploy a variety of strategies to deprive access of invading pathogens to trace metals; an active process known as nutritional immunity (2, 5–8). To date, the best characterized nutritional immunity strategy is based on mobilization of metal-chelating proteins to the infection site by a variety of host immune cells (2). Among them, calprotectin, a member of the S100 protein family produced by neutrophils and other types of immune cells that is secreted in large quantities during infection and inflammatory processes, is the main host protein responsible for Mn^2+^ and Zn^2+^ sequestration (9–11). To overcome trace metal limitation, microbial pathogens evolved effective metal-scavenging systems that include expression of cell surface-associated high-affinity metal uptake transporters; and in some bacterial species, synthesis and trafficking of organic extracellular molecules known as metallophores (8).

Although the importance of Fe in host-pathogen interactions has been heavily examined (6), the role of Mn and Zn in host-pathogen interactions and the mechanisms utilized by bacteria to maintain their cellular levels and ratios properly balanced are less understood (7, 12–17). The second most abundant trace metal in vertebrates, Zn is estimated to be incorporated into approximately 5% of the bacterial proteome serving structural and catalytic roles in multiple biological processes (18, 19). In most bacterial pathogens, Zn homeostasis under the severe Zn restricted conditions that can be encountered in host environments depends on the activity of surface-associated Zn uptake systems from the ATP-binding cassette (ABC) transporter family (reviewed in (20–22)). In addition, important human pathogens such as *Pseudomonas aeruginosa* and *Staphylococcus aureus* produce Zn-binding metallophores, also known as zincophores (23).

To date, the contributions of Zn acquisition systems to virulence has been demonstrated in a number of bacterial species, including several Gram-positive pathogens that are phylogenetically-related to *Enterococcus faecalis*, the subject organism of the present study. In *S. aureus*, inactivation of either the ABC-type transporter AdcABC, the staphylopine (Stp) zincophore, or its cognate multi-metal transporter CntABCDF was sufficient to impair bacterial growth under Zn restricted conditions *in vitro* (24). Loss of both AdcABC and Stp/CntABCDF systems resulted in further growth impairment under Zn restricted conditions and attenuated virulence in a mouse retro-orbital infection model (24). In streptococci, which to date reportedly do not synthesize zincophores, Zn acquisition is primarily mediated by the ABC-type transporter AdcABC. In addition to AdcABC, most streptococcal species encode an additional *adcA* homologue, known as *adcAII* (reviewed in (25)), coding for a second Zn-binding lipoprotein. In *Streptococcus pyogenes*, strains lacking *adcC*, *adcA* or *adcAII* grew poorly in the presence of purified human calprotectin and displayed attenuated virulence in a necrotizing fasciitis mouse model (26) and in a humanized-plasminogen skin infection mouse model (27). Notably, *S. pyogenes* Δ*adc* strains were shown to retain wild-type strain levels of virulence in calprotectin-negative (*S100a9*^-/-^) mice (26), which validates the central role of calprotectin in host-mediated Zn sequestration and protection against bacterial infection. Similarly, virulence of *Streptococcus pneumoniae* Δ*adcA*Δ*adcAII* and *Streptococcus agalactiae* Δ*adcA*Δ*adcAII*Δ*lmb* (*lmb* encodes for a 3^rd^ Zn-binding lipoprotein) mutants were significantly attenuated in mouse models of systemic and nasopharyngeal colonization (16, 28). Finally, in the oral pathogen *S. mutans,* one of the few streptococci that do not encode the orphan *adcAII* gene, inactivation of the *adcABC* system halted bacterial growth under Zn-restricted conditions and severely impaired bacterial colonization of the dental biofilm in a rat model (12, 29).

A commensal of the gastrointestinal (GI) tract, *E. faecalis,* is also a prevalent opportunistic pathogen of localized and systemic infections, including but not limited to infective endocarditis, catheter-associated urinary tract infections (CAUTI), and wound infections (30–33). A major virulence trait of *E. faecalis* is its remarkable capacity to adapt to adverse conditions in the GI tract (their natural host environment) and several other host tissues, and to resist (or tolerate) exposure to hospital-grade disinfectants and antibiotic treatments (34, 35). Because very little is known about the mechanisms of Zn homeostasis in enterococci, we sought to characterize the Zn acquisition systems of *E. faecalis* in this study. Similar to most streptococci, the genome of *E. faecalis* encodes for a conserved AdcACB system (originally annotated as *znuACB*) and an orphan substrate-binding lipoprotein AdcA-II that has been annotated as *adcA*. In this report, we isolated a panel of *E. faecalis* Δ*adc* strains, including strains lacking every *adc* gene (Δ*adcACBΔadcAII*) or both genes coding for the substrate-binding lipoproteins (Δ*adcAΔadcAII*), and then used these mutants to assess the role of AdcACB and AdcAII in *E. faecalis* pathophysiology. Our results revealed that simultaneous inactivation of *adcA* and *adcAII* or of the entire *adcACB* operon yielded the most impactful phenotypes, which included severe growth/survival defects in the presence of calprotectin or in human serum *ex vivo*, and attenuated virulence in both invertebrate and vertebrate infection models. We also discovered that the inability to maintain Zn homeostasis diminished the notorious high tolerance of *E. faecalis* to antibiotics that target the cell envelope, which may be linked to alterations in fatty acid species profile and abundance. Collectively, this study reveals that AdcACB and AdcAII work cooperatively to maintain *E. faecalis* Zn homeostasis during infection such that the surface-associated AdcA and AdcAII lipoproteins can be considered as potential targets for the development of antimicrobial interventions.

## RESULTS

### AdcACB and AdcAII work in concert to promote growth under Zn-restricted conditions

Using the PubMed BLASTn search tool, we identified the genes coding for the conserved ABC-type transporter AdcACB (*OG1RF_RS00260-RS00270*) and the orphan substrate-binding AdcAII protein (*OG1RF_RS12625*) (**Fig. 1A**) in the *Enterococcus faecalis* OG1RF genome (GenBank: CP002621.1). The translated gene products of *OG1RF_RS00260* (Accession ID: AEA92738.1) and *OG1RF_RS12625* (Accession ID: AEA95159.1) display 57% and 64% amino acid similarity to, respectively, the Zn-binding lipoprotein AdcA and AdcAII of *S. pneumoniae* R6 (16). Pairwise alignment between the translated gene products of *OG1RF_RS00260* and *OG1RF_RS12625* revealed 53% similarity, suggesting functional redundancy between AdcA and AdcAII. Furthermore, the translated gene products of *OG1RF_RS00265* and *OG1RF_RS00270* displayed 61.67% and 55.47% similarity to, respectively, the *S. mutans* AdcB (membrane permease) and AdcC (cytoplasmic ATPase) (29). In prior transcriptome-based studies conducted with *E. faecalis* strain V583, the *adcABC* (originally annotated as *znuABC*) and *adcAII* genes were shown to be repressed after exposure to high Zn levels and strongly induced after treatment with the Zn-chelating agent TPEN (N,N,N′,N′-tetrakis(2-pyridinylmethyl)-1,2-ethanediamine) (36, 37). Based on the presence of conserved domains and amino acid similarities with homologous systems of closely-related streptococci, we renamed the *OG1RF_RS00260-OG1RF_RS00270* gene cluster *adcACB*, and used the *adcAII* designation for the lone *OG1RF_RS12625*. Of note, none of the enterococcal genomes surveyed, including *E. faecalis* OG1RF, encode biosynthetic gene clusters and cognate transporters of opine-like zincophore systems that have been identified in a small number of bacterial pathogens (23).

**Fig 1.**
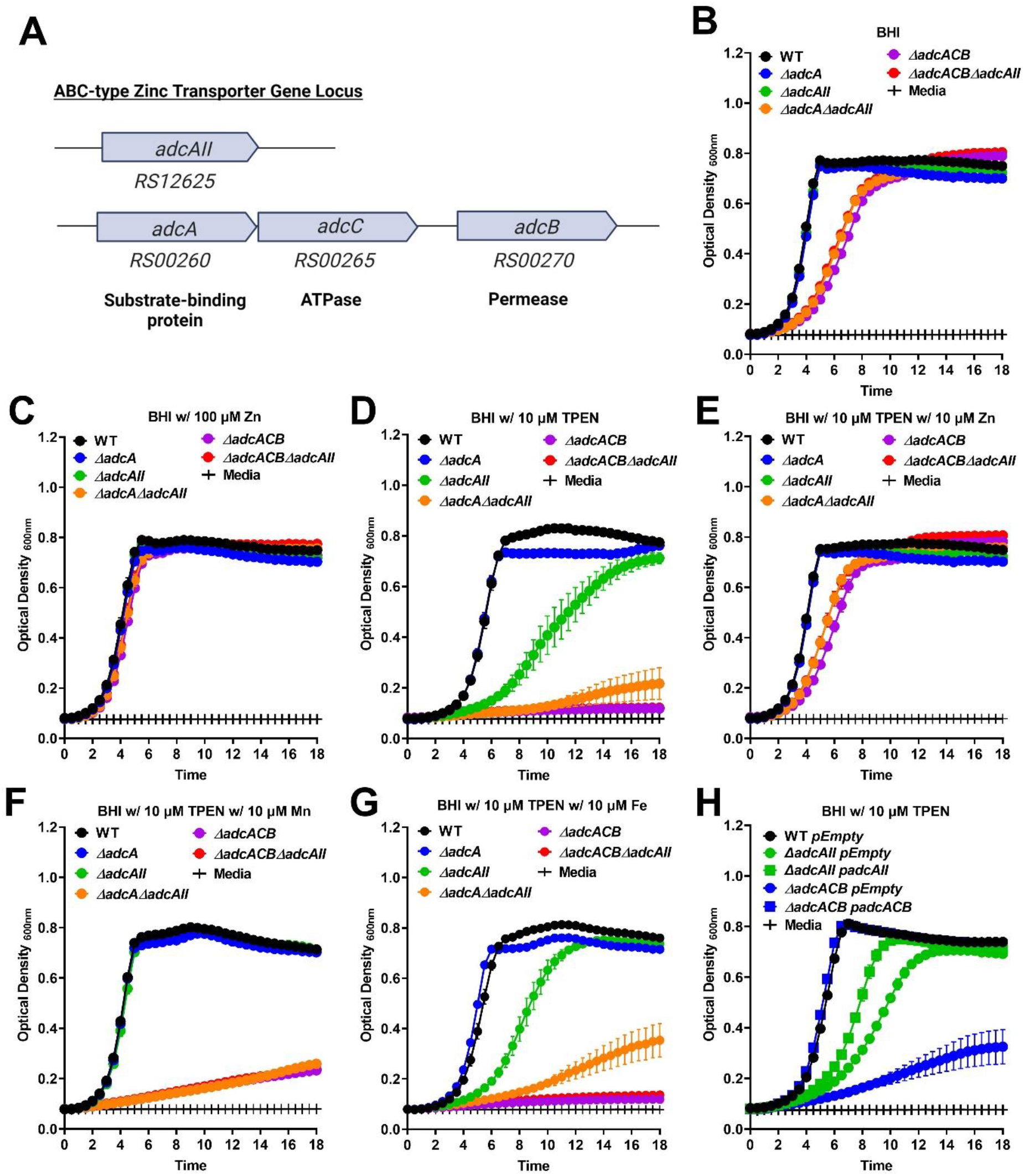
Growth characteristics of *E. faecalis* and its Zn-deficient mutants under Zn-restricted conditions. (A) Schematic of the gene locus of the Zn transporters in *E. faecalis* OG1RF core genome. Growth curves of *E. faecalis* wild type (WT) and *Δadc* mutants in BHI (B), BHI supplemented with 100 µM ZnSO_4_ (C), 10 µM TPEN (D) or combination of TPEN and ZnSO_4_ (E), TPEN and MnSO_4_ (F), or TPEN and Fe SO_4_ (G). For B, D-F, data points represent the average of nine biological replicates. Lastly, growth curve of *E. faecalis* wild type (WT) and genetically complemented Δ*adc* mutants in BHI supplemented with 10 µM TPEN (H). For C and G, data points represent the average of six biological replicates. Error bar represent the standard error of margin (SEM). Statistical analysis was performed using simple linear regression of exponential growth phase, and slope of each mutant’s growth kinetics was compared to the parent strain.

To probe the role of AdcACB and AdcAII in Zn acquisition and, most importantly, determine the significance of these systems to *E. faecalis* pathophysiology, we used a markerless in-frame deletion strategy (38) to isolate strains lacking one or both substrate-binding lipoproteins (*ΔadcA, ΔadcAII*, *ΔadcAΔadcAII*), the entire *adcACB* operon alone or in combination with *adcAII* (Δ*adcACB* and Δ*adcACBΔadcAII*). Then, we compared the ability of *E. faecalis* OG1RF (wild-type strain) and Δ*adc* derivatives to grow in BHI, a complex media that contains ∼10 µM Zn (39), or BHI supplemented with the Zn-specific chelator TPEN (40). In BHI, inactivation of either *adcA* or *adcAII* alone had no impact in growth kinetics or growth rates whereas simultaneous inactivation of both Zn-binding lipoproteins (*ΔadcAΔadcAII*) or entire *adcACB* operon (Δ*adcACB* and Δ*adcACBΔadcAII*) resulted in slower growth rates that did not compromise final growth yields (**Fig. 1B**). Supplementation of BHI with 100 µM ZnSO_4_ restored the growth defects of Δ*adcAΔadcAII*, Δ*adcACB* and Δ*adcACBΔadcAII* mutants (**Fig. 1C**). While addition of 10 µM TPEN to BHI (BHI+TPEN) had very little impact on growth of the parent OG1RF strain (OG1RF growth is only severely impaired at 20 µM TPEN, **Fig. S1**), it completely inhibited growth of Δ*adcACB* and Δ*adcACBΔadcAII*, while also inhibiting the growth of *ΔadcAII* (**Fig. 1D**); by contrast, addition of 10 µM ZnSO_4_ to BHI+TPEN alleviated TPEN-mediated Zn restriction effects on these mutants (**Fig. 1E**). To confirm that the inhibitory effect of TPEN is Zn-specific, we also tested whether addition of 10 µM MnSO_4_ or 10 µM FeSO_4_ could restore growth of the Δ*adc* strains. Excluding the observation that Mn supplementation restoring the growth defect of *ΔadcAII*; restoration of growth defects was not observed in all other strains regardless of Fe or Mn supplementation (**Fig. 1F-G**, compare to **Fig. 1D**). Finally, *in trans* complementation of Δ*adcACB* and of *ΔadcAII*, respectively, fully or partially rescued their growth defects in BHI+TPEN (**Fig. 1H**). The reasons for manganese rescuing grown of *ΔadcAII* in the presence of TPEN and the partial complementation of *ΔadcAII* are presently unknown.

Next, we sought to determine the fitness of our panel of Δ*adc* strains to compete with the divalent metal-sequestering host metalloprotein calprotectin, which we previously showed is a potent inhibitor of *E. faecalis* strains that lack two or more of its three major Mn import systems (41). We carried out growth kinetic assays to compare the ability of OG1RF (WT) and Δ*adc* mutants to grow in BHI supplemented with wild-type human calprotectin (hCP) or recombinant calprotectin (hCPΔ_Mn-tail_) defective in Mn sequestration (gifts from Dr Walter Chazin, Vanderbilt University, USA)(10). In the presence of hCP, growth of *ΔadcA* was not significantly different when compared to WT whereas Δ*adcAII,* Δ*adcA*Δ*adcAII,* Δ*adcACB* and Δ*adcACB*Δ*adcAII* mutants displayed reduced growth rates (**Fig. 2A**). While inactivation of the manganese (Mn)-binding residue in hCP_ΔMn-tail_ improved growth rates of parent and *ΔadcA* strains, all other mutants remained highly sensitive to hCP_ΔMn-tail_ displaying reduced growth rates (**Fig. 2B**, compare with panel **2A**). *In trans* complementation fully rescued the growth defects of Δ*adcACB* mutant in the presence of both versions of calprotectin, whereas Δ*adcAII* in hCP_ΔMn-tail_ only (**Fig. 2C-D**). The reason for the partial complementation of Δ*adcAII* in the presence of hCP is presently unknown.

**Fig 2.**
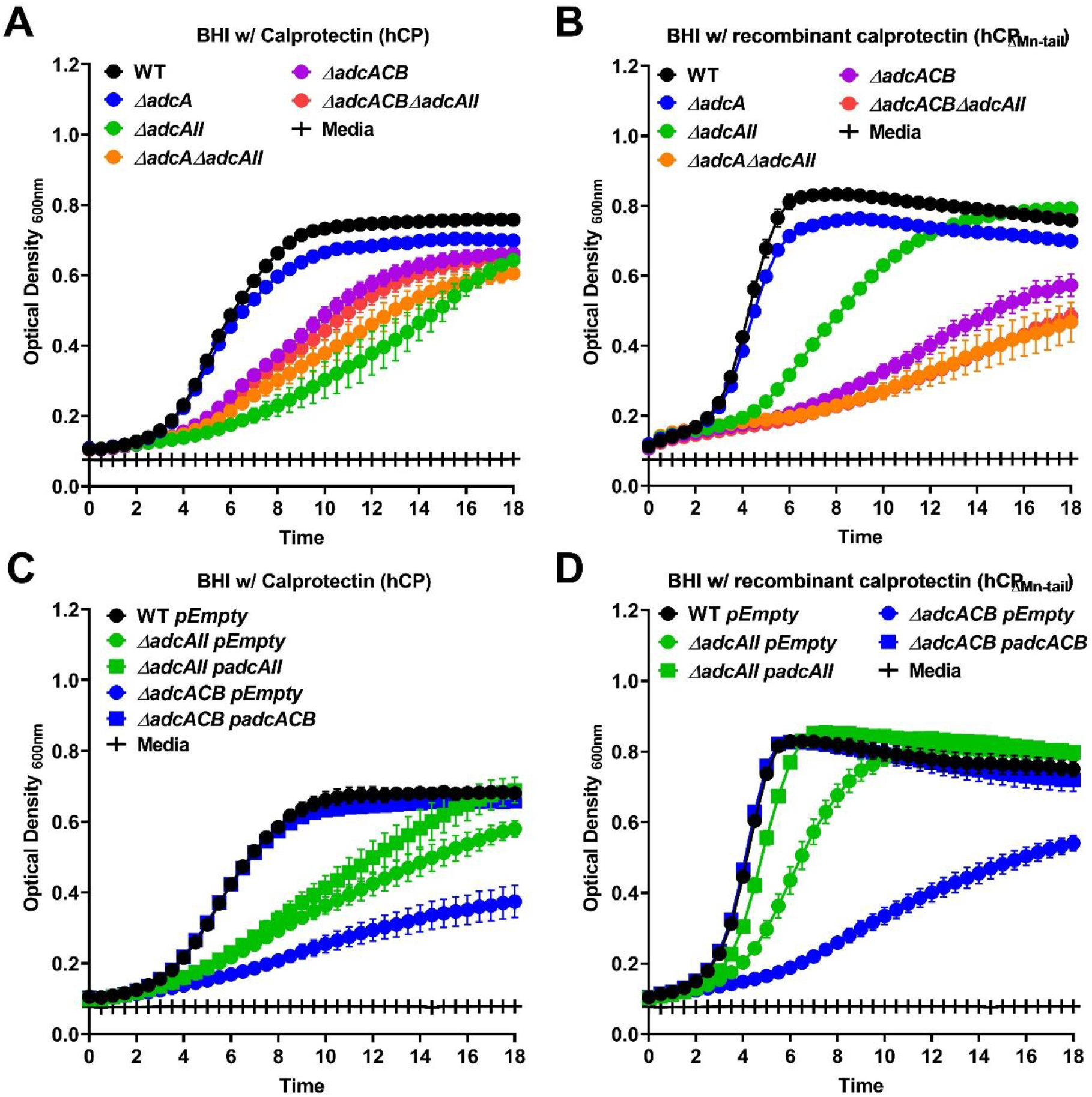
Growth characteristics of *E. faecalis* and its Zn-deficient mutants in the presence of calprotectin. Growth curves of *E. faecalis* OG1RF WT and its mutants (A-B), as well as the genetic complemented Zn-deficient mutants (C-D), in BHI supplemented with CP buffer and 150 µg ml^-1^ of WT hCP or hCP_ΔMn-tail_. For A-B and C-D, data points represent the average and error bar represent the standard error of margin (SEM) of nine and six biological replicates, respectively. Statistical analysis was performed using simple linear regression of exponential growth phase, and slope of each mutant’s growth kinetics was compared to the parent strain.\

Taken together, these findings reveal that the lipoproteins AdcABC and AdcAII work independently but cooperatively to facilitate *E. faecalis* growth under Zn-restricted conditions. Based on the identical phenotypes of Δ*adcACB* and *ΔadcAΔadcAII* strains, these results also (strongly) suggest that both AdcA and AdcAII associate with AdcB (inner membrane permease) and AdcC (cytoplasmic ATPase) to form tripartite Zn transporters. Finally, growth kinetics in the presence of Zn-chelating agents hint that AdcAII is more effective than AdcA at mediating Zn acquistion under the more severely Zn-restricted conditions.

Next, we used inductively-coupled optical emission spectrometry (ICP-OES) to determine intracellular Zn pools in mid-log grown cultures of WT and derivative Δ*adc* strains grown in BHI or BHI supplemented with 7.5 µM TPEN (**Fig. 3A**). Of note, this concentration of TPEN is sub-inhibitory to the growth of Δ*adcACB* and of *ΔadcAΔadcAII* (data not shown), such that all cell cultures could be harvested at the OD_600_ of 0.5. In BHI, only Δ*adcACB* was found to accumulate statistically significantly less Zn intracellularly when compared to the WT strain albeit this difference was rather small (∼12%), and not observed in Δ*adcA*Δ*adcAII* and Δ*adcACB*Δ*adcAII* strains (**Fig. 3A**), possibly due to the large variability among biological replicates in these mutants. The addition of TPEN to the growth media led to a significant, and unexpected, increase in intracellular Zn pools in the WT strain when compared to cells grown in BHI (∼ 50% increase). Nonetheless, all mutants accumulated less Zn compared to the WT strain grown in BHI+TPEN whereas only Δ*adcACB*Δ*adcAII* mutant accumulated significantly less (> 50% decrease) intracellular Zn as compared to cells grown in BHI (**Fig. 3A**). We suspected that the higher intracellular levels of Zn of the parent strain grown in BHI+TPEN correlated with increased transcription of the *adcACB* and *adcAII* genes. To verify this possibility, we used quantitative RT-PCR to determine mRNA levels of *adcA* and *adcAII* in the WT strain grown to mid-log phase in BHI and then treated with 30 µM TPEN or 4 mM ZnSO_4_ for 1 hour. As anticipated, and in line with previous transcriptional studies (36, 37), TPEN treatment increased transcript levels of *adcA* (∼1-log) and *adcAII* (∼2-log) whereas Zn supplementation reduced mRNA levels by ∼2-log (*adcA*) and ∼1-log (*adcAII*) respectively, when compared to untreated (BHI) control (**Fig. 3B**). Thus, it appears that while TPEN chelates Zn, the upregulation of *adc* genes allows *E. faecalis* to overcome Zn limitation, at least to a certain point.

**Fig 3.**
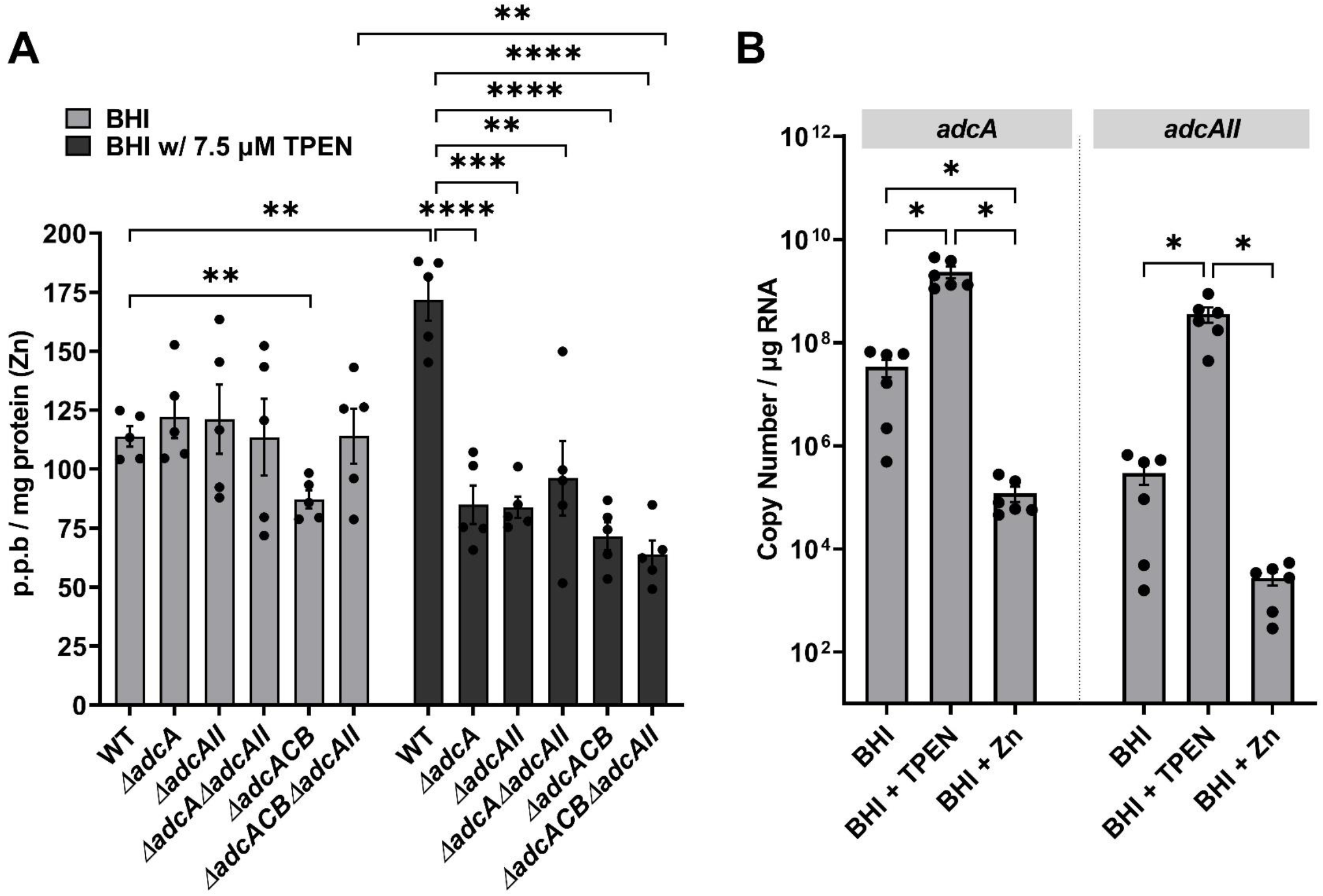
Intracellular Zn quantification and transcriptional profiles of *E. faecalis* and its Zn-deficient mutants. (A) ICP-OES quantifications of intracellular Zn of mid-log grown *E. faecalis* OG1RF WT and derivatives grown in BHI orand BHI supplemented with 7.5 µM TPEN. Data points represent five biological replicates. Statistical analysis was performed using two-way ANOVA with Dunnett’s multiple comparison test. ** *p* ≤ 0.01, *** *p* ≤ 0.001 and **** *p* ≤ 0.0001. (B) Comparison of reversed transcribed cDNA copy of *adcA* and *adcAII* in *E. faecalis* OG1RF grown for 1 hour in BHI, BHI with 30 µM TPEN and BHI with 4 mM ZnSO_4_. Data points represent the average of six biological replicates. Statistical analysis was performed using unpaired *t*-test with Welch’s correction. * *p* ≤ 0.05. Error bar represent the standard error of margin (SEM).

### The lower intracellular Zn levels of *Δadc* strains is accompanied by reduction in Mn pools

In addition to Zn, we determined intracellular levels of Mn in the *Δadc* strains grown in BHI or BHI+TPEN. While there were no significant differences in Mn pools of strains grown in BHI, all mutants accumulated significantly less (∼50% reduction) Mn when compared to the WT strain (**Fig. S2A**). This result was somewhat unexpected considering that *adc* gene deletion in both *S. mutans* and *S. pneumoniae* led to Mn accumulation under Zn-restricted conditions (2, 16, 29). Because a balanced Mn:Zn ratios is critical to cell homeostasis, we speculate that lowering Mn uptake/accumulation is an adaptive mechanism of *E. faecalis* Δ*adc* strains to maintain cell homeostasis. To explore this possibility, we tested the ability of WT and Δ*adcACBΔadcAII* strains to grow in BHI containing increasing concentrations of MnSO_4_. While growth of WT was not affected by increasing Mn concentrations (up to 0.5 M MnSO_4_), we observed that Δ*adcACBΔadcAII* mutant displayed significant growth delay as compared to the WT strain (**Fig. S2B-C**).

### AdcACB and AdcAII contribute to growth in serum but not in urine *ex vivo*

To determine the contribution of Adc-mediated Zn uptake to *E. faecalis* virulence, we first monitored the ability of the Δ*adc* strains to grow and survive in pooled human serum or human urine *ex vivo*. When incubated in serum, growth of Δ*adcA* and Δ*adcAII* strains did not significantly differ from WT whereas Δ*adcA*Δ*adcA,* Δ*adcACB* and Δ*adcACB*Δ*adcA* grew poorly and, most relevantly, displayed sharp decreases in survival after 8 hours and onwards; ultimately showing ∼3-log reduction in colony forming unit (CFU) recovered after 48 hours of incubation in serum (**Fig. 4A**). These growth and survival defects were fully reversed by Zn (500 µM ZnSO_4_) supplementation (**Fig. 4B**) or by genetic complementation (**Fig. 4C**). On the other hand, the ability of all Δ*adc* strains to grow/ survive in urine was not found to differ from WT (**Fig. 4D**), suggesting that Zn is not a growth-limiting factor in urine. While total or free Zn levels in serum or in urine were not determined in this study, the strong phenotype of the Δ*adcA*Δ*adcAII,* Δ*adcACB* and Δ*adcACB*Δ*adcA* strains in serum and lack of phenotype in urine are not completely unexpected. In circulation, total Zn levels is estimated to be low with most Zn sources bound or sequestered to host cells and proteins (42, 43). On the other hand, Zn is abundant in the bladder environment as any excess Zn, typically from dietary sources, is excreted through urine via the gastrointestinal route (44–47).

**Fig 4.**
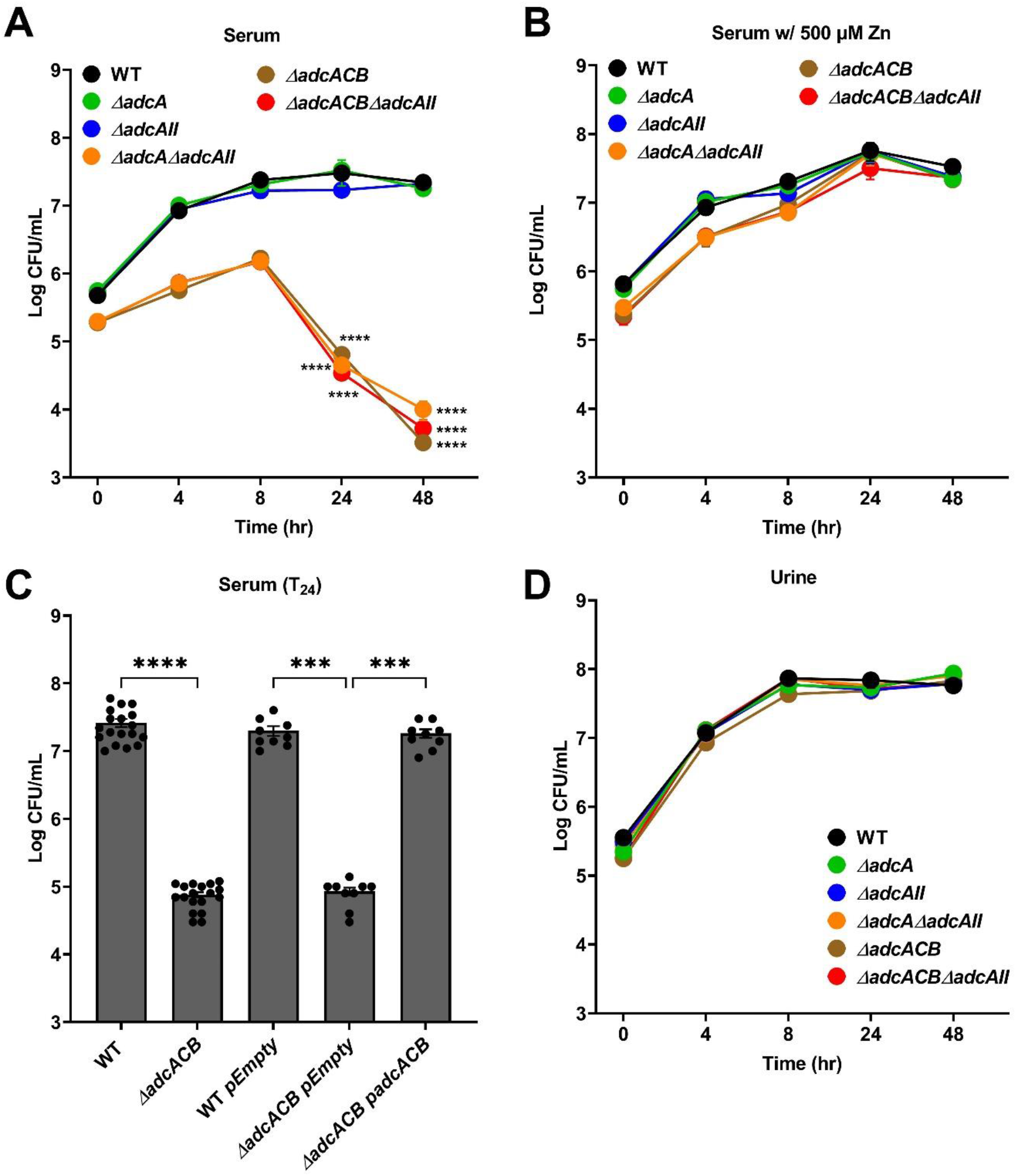
Growth and survival of *E. faecalis* in serum and urine. Colony-forming units (CFU) of *E. faecalis* OG1RF WT and its mutants incubated in pooled (A) human serum or (B) serum supplemented with 500 µM ZnSO_4_, and pooled human urine (D). (C) CFU counts of *E. faecalis* OG1RF WT, its mutants and genetic complemented mutants after 24 hours incubation in pooled human serum. For A-D, data points represent the average and error bar represent the standard error of margin (SEM) of nine biological replicates. Statistical analysis was performed using one-way ANOVA with Welch’s correction. *** *p* ≤ 0.001 and **** *p* ≤ 0.0001.

### Disruption of AdcABC-AdcAII lowers tolerance towards cell envelope-targeting antibiotics

Because the Δ*adcA*Δ*adcAII,* Δ*adcACB* and Δ*adcACB*Δ*adcAII* strains displayed a tendency to clump in liquid culture (data not shown), we wondered if this was caused by altered chain length formation. Indeed, these mutants formed longer chains when compared to the WT, Δ*adcA* and Δ*adcAII* strains that primarily formed very short chains or diplococcus (**Fig. 5**). This observation and the fact that an *S. pneumoniae* Δ*adcA*Δ*adcAII* mutant strain displayed aberrant cell septation (16), led us to wonder if expression of virulence traits that occur at the cell surface interface were affected in the mutant strains. First, we compared the capacity of WT and mutants to form biofilms after 24 hours incubation in BHI supplemented with 10 mM glucose. The total biofilm biomass of Δ*adcA* and Δ*adcAII* single mutants was significantly reduced when compared to WT, but the very small differences observed (5 to 10% reduction) are unlikely to have major biological implications (**Fig. 6A**). On the other hand, the Δ*adcA*Δ*adcAII,* Δ*adcACB* and Δ*adcACB*Δ*adcAII* strains formed more robust biofilms with ∼30 to 50% increase in biofilm biomass (**Fig. 6A**).

**Fig 5.**
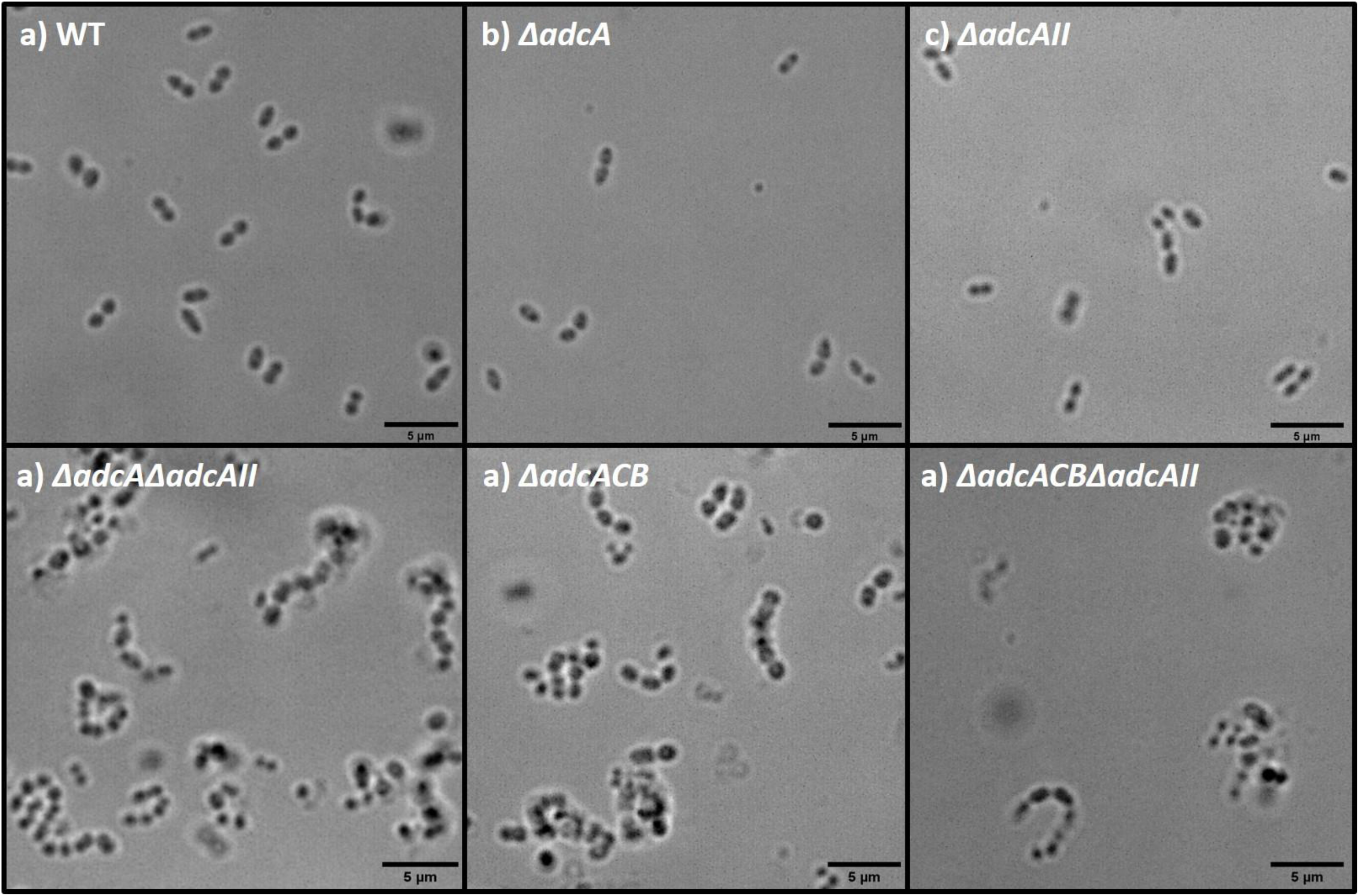
Bright-field microscopic images of *E. faecalis* and its indicated mutants. Images shown are representative of 10 images that are acquired from one biological sample from each strain grown in BHI respectively. Black bar represent five microns in length.

**Fig 6.**
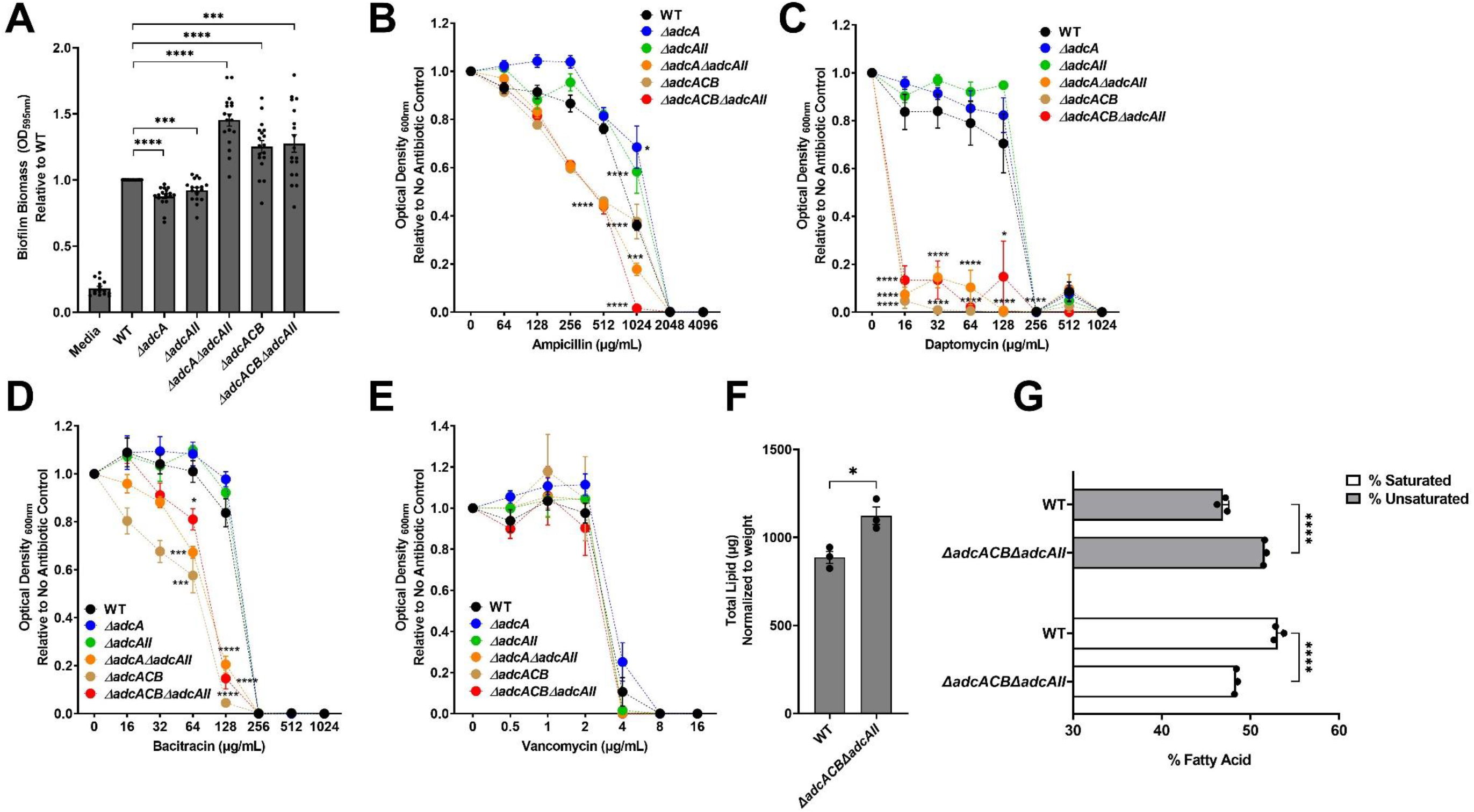
Characterization of *E. faecalis* virulence traits at cell surface interface. (A) Biofilm biomass quantification of *E. faecalis* OG1RF WT and its mutants grown in BHI for 24 hours. Statistical analysis was performed using One-way ANOVA with Welch’s correction. (B-E) Final growth yields of *E. faecalis* OG1RF WT and its mutants after 24 hours incubation in BHI supplemented with 2-fold increasing concentrations of (B) ampicillin, (C) daptomycin, (D) bacitracin and (E) vancomycin. For A-E, data points represent the average of nine biological replicates. Statistical analysis was performed using One-way ANOVA with Welch’s correction. * *p* ≤ 0.05, *** *p* ≤ 0.001 and **** *p* ≤ 0.0001. For A-E, error bars represent the standard error of margin (SEM). (F-G) Gas chromatography fatty acid methyl ester (GC-FAME) analysis of total lipid content (F) and saturated/unsaturated fatty acids percentages (G) of cell lysates harvested from *E. faecalis* OG1RF and *ΔadcACBΔadcA-II* mutant. Data points represent the average of three biological replicates. Statistical analysis was performed using unpaired student t-test. * *p* ≤ 0.05, ** *p* ≤ 0.01, *** *p* ≤ 0.001 and **** *p* ≤ 0.0001. For F-G, error bars represent the standard deviation (SD).

Next, we tested the capacity of WT and mutants to grow in the presence of antibiotics that target different steps of cell wall biosynthesis (ampicillin, bacitracin and vancomycin) or disrupt membrane integrity (daptomycin) by determining the minimal inhibitory concentration (MIC) for each antibiotic. Once again, relevant phenotypes were restricted to the Δ*adcA*Δ*adcAII,* Δ*adcACB* and Δ*adcACB*Δ*adcAII* strains. Specifically, Δ*adcA*Δ*adcAII,* Δ*adcACB* and Δ*adcACB*Δ*adcAII* showed lower MICs for ampicillin, bacitracin and daptomycin (**Fig. 6B-D**). However, WT and all mutants showed the same vancomycin MIC (**Fig. 6E**). Due to the extremely high daptomycin sensitivity of some of the mutants (WT MIC = 256 μg ml^-1^, Δ*adcA*Δ*adcAII,* Δ*adcACB* and Δ*adcACB*Δ*adcAII* MIC = 16 μg ml^-1^ml) and knowing that daptomycin tolerance of *E. faecalis* is, in great part, dependent on changes in membrane fatty acids profile (48–50), we used gas chromatography analysis of fatty acid methyl esters (GC-FAME) to determine the fatty acid profile of WT and Δ*adcACB*Δ*adcAII* strains grown in BHI. Of the 21 fatty acid species detected, the abundance of 10 fatty acids was significantly altered in Δ*adcACB*Δ*adcAII* when compared to WT (**Table 1**). Most notably, two unsaturated fatty acids, oleic acid and linoleic acid, associated with daptomycin tolerance (48) were significantly more abundant in the mutant strain. Moreover, the Δ*adcACB*Δ*adcAII* strain displayed higher total lipid content with a slightly higher ratio of unsaturated fatty acids over saturated fatty acids while the WT strain displayed greater abundance of saturated fatty acids than unsaturated fatty acids (**Fig. 6F-G**).

**Table 1.** GC-FAME analysis.

| Mean value ± standard deviation (µg lipid/cell weight) |  |  |  |  |  |  |
| --- | --- | --- | --- | --- | --- | --- |
| Fatty Acid Group | Common Name | Speciation | *E. faecalis* WT | Δ*adcACB*Δ*adcA-II* | P-value | Significance |
| Saturated | Caprylic acid | **C8:0** | 0.065 ± 0.019 | ND | - | - |
| Saturated | Capric acid | **C10:0** | 0.827 ± 0.121 | 0.617 ± 0.059 | 0.0758 | * |
| Saturated | Lauric acid | **C12:0** | 7.310 ± 0.985 | 7.920 ± 0.297 | 0.3982 | n.s |
| Saturated | Myristic acid | **C14:0** | 69.780 ± 9.570 | 61.557 ± 3.604 | 0.2724 | n.s |
| Mono-unsaturated | Myristoleic acid | **C14:1n9** | 1.167 ± 0.254 | 1.453 ± 0.215 | 0.2118 | n.s |
| Saturated | Pentadecylic acid | **C15:0** | 0.757 ± 0.153 | 1.307 ± 0.231 | 0.0328 | * |
| Saturated | Palmitic acid | **C16:0** | 358.023 ± 15.292 | 432.55 ± 30.600 | 0.0337 | * |
| Mono-unsaturated | Palmitoleic acid | **C16:1n9** | 89.840 ± 12.827 | 107.423 ± 5.350 | 0.1269 | n.s |
| Saturated | Margaric acid | **C17:0** | 0.417 ± 0.104 | 0.750 ± 0.173 | 0.0583 | n.s |
| Saturated | Stearic acid | **C18:0** | 33.180 ± 4.151 | 38.540 ± 5.094 | 0.2333 | n.s |
| Mono-unsaturated | Elaidic acid | **C18:1n9** | 317.087 ± 22.463 | 456.567 ± 37.721 | 0.0095 | ** |
| Di-unsaturated | Linoleic acid | **C18:2n6** | 2.530 ± 0.682 | 4.770 ± 0.993 | 0.0384 | * |
| Di-unsaturated | gamma-Linolenic acid | **C18:3n6** | 0.059 ± 0.011 | 0.127 ± 0.021 | 0.0153 | * |
| Di-unsaturated | α-linolenic acid | **C18:3n3** | 0.893 ± 0.248 | 1.477 ± 0.341 | 0.0807 | n.s |
| Saturated | Arachidic acid | **C20:0** | 0.217 ± 0.035 | 0.313 ± 0.076 | 0.1443 | n.s |
| Mono-unsaturated | 11-Eicosenoic acid | **C20:1n9** | 0.513 ± 0.119 | 1.090 ± 0.257 | 0.0427 | * |
| Mono-unsaturated | Eicosadienoic acid | **C20:2n6** | 0.363 ± 0.042 | 0.383 ± 0.060 | 0.6639 | n.s |
| Di-unsaturated | Dihomo-gamma-linolenic acid | **C20:3n6** | ND | 0.110 ± 0.017 | - | - |
| Tetra-unsaturated | Arachidonic acid | **C20:4n6** | 0.850 ± 0.233 | 1.697 ± 0.299 | 0.0202 | * |
| Tetra-unsaturated | cis-11,14,17-Eicosatrienoic acid | **C20:3n3** | ND | 0.068 ± 0.009 | - | - |
| Penta-unsaturated | Eicosapentaenoic acid | **C20:5n3** | 0.072 ± 0.014 | 0.120 ± 0.017 | 0.0219 | * |
| Saturated | Behenic acid | **C22:0** | ND | 0.078 ± 0.015 | - | - |
| Mono-unsaturated | Erucic acid | **C22:1n9** | 2.530 ± 0.682 | 4.770 ± 0.993 | 0.0384 | * |
| Saturated | Tricosylic acid | **C23:0** | 0.057 ± 0.005 | ND | - | - |
| Tetra-unsaturated | docosatetraenoate | **C22:4n6** | 0.363 ± 0.552 | 0.084 ± 0.015 | 0.4741 | n.s |
| Penta-unsaturated | Docosapentaenoic acid | **C22:5n3** | ND | 0.107 ± 0.019 | - | - |
| Hexa-unsaturated | Docosahexaenoic acid | **C22:6n3** | 0.110 ± 0.040 | 0.103 ± 0.015 | 0.8076 | n.s |

### Zn is critical for *E. faecalis* virulence in multiple animal infection models

In the last series of experiments, we used three *in vivo* models to probe the contributions of AdcABC and AdcAII to virulence. In conformity with *in vitro* and *ex vivo* phenotypes, virulence of Δ*adcA*Δ*adcAII,* Δ*adcACB* and Δ*adcACB*Δ*adcAII* but not the single mutants Δ*adcA* and Δ*adcAII* was highly attenuated in the *Galleria mellonella* model (**Fig. 7A**). Next, we used two catheter-associated mouse infection models that recapitulate some of the environmental and immunological conditions that promote enterococcal infections in human. From this point, we did not include the Δ*adcA*Δ*adcAII* and Δ*adcACB* strains as these mutants always phenocopied the Δ*adcACB*Δ*adcAII* mutant. In the catheter-associated peritonitis model, the Δ*adcA* and Δ*adcAII* single mutants colonized peritoneal cavity, catheter and spleen (infection becomes systemic after 12 to 24 h) in significantly fewer numbers than the WT strain (**Fig. 7B**). Not surprisingly, colonization defects of single mutants were significantly more pronounced in the Δ*adcACB*Δ*adcAII* strain. Lastly, in a catheter-associated urinary tract infection (CAUTI) model, virulence of Δ*adcA* and Δ*adcACB*Δ*adcAII* strains was attenuated but bacterial burden recovered from bladders of retrieved catheter of animals infected with WT or Δ*adcAII* strains were nearly identical (**Fig. 7C**). Moreover, Δ*adcAII* was recovered from kidneys and spleen in higher bacterial titers, albeit the trend of the later was not considered significant (**Fig. 7C**).

**Fig 7.**
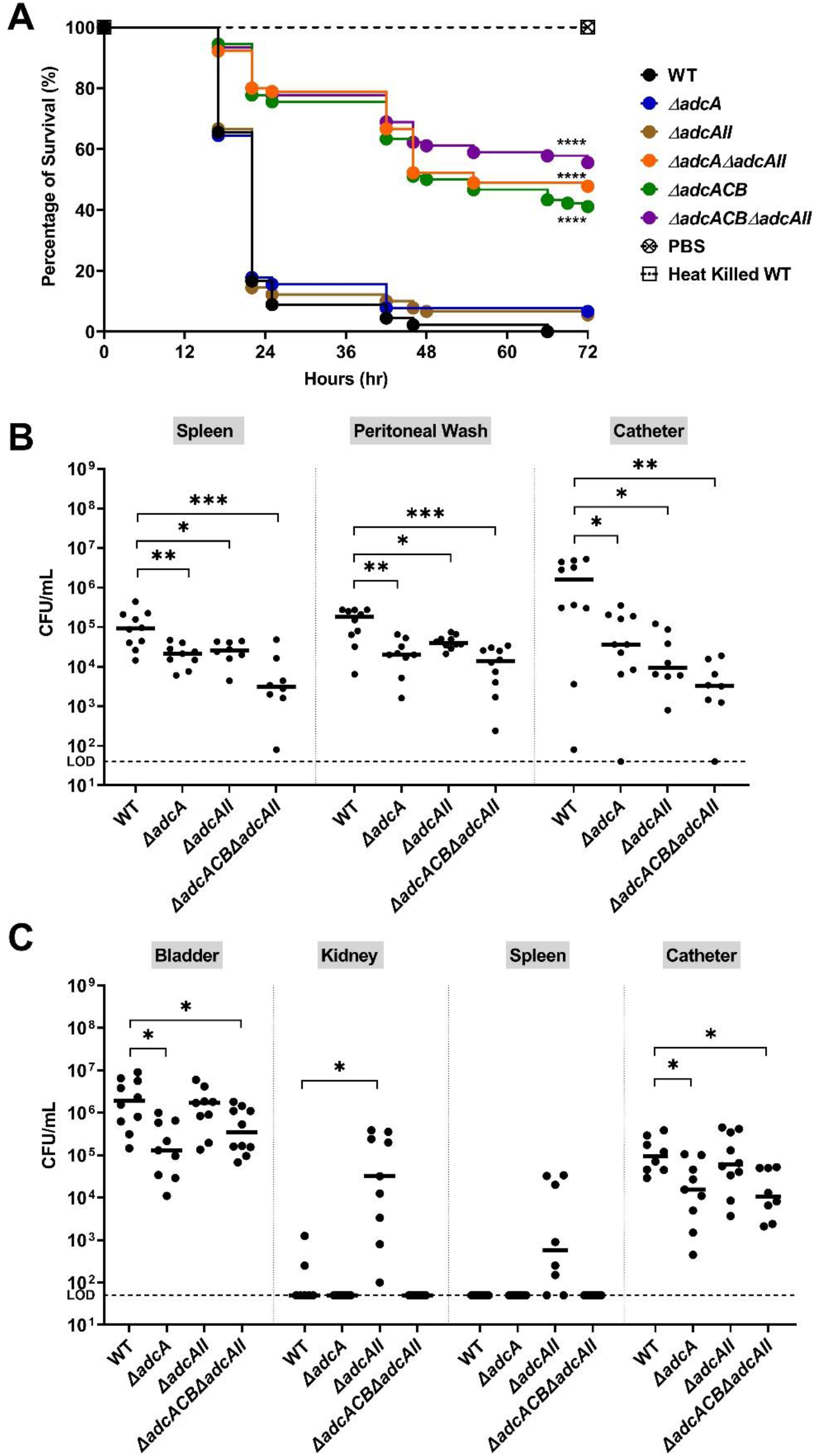
Virulence of *E. faecalis* in different animal models. (A) Percentage survival of *G. mellonella* larvae 96 hours post-infection with *E. faecalis* WT or indicated mutants. Each curve represents a group of 15 larvae injected with ∼1 x 10^5^ CFU of selected *E. faecalis* strain. Data points represent average of 6 biological replicates. Statistical analysis was performed using Log-rank (Mantel-Cox) test. (B) Total CFU recovered after 48 hours from spleen, peritoneal wash and catheter of mice infected with 2 x 10^8^ CFU of bacteria. (C) Total CFU recovered after 24 hours from bladder, kidney, spleen and catheter of mice infected with 1 x10^7^ CFU of WT or indicated mutants. For (B-C), ten mice were infected with two biological replicates and data points shown were a result of using ROUT outlier test. Black line represents the median. Statistical analysis was performed using Mann-Whitney test. * *p* ≤ 0.05, ** *p* ≤ 0.01, *** *p* ≤ 0.001, **** *p* ≤ 0.0001. The dashed line represents the limit of detection (LOD = 50 CFUs).

## DISCUSSION

While there has been increasing appreciation of the multiple contributions of trace metals other than Fe to bacterial fitness and virulence (2, 5, 20), the mechanisms utilized by *E. faecalis* to maintain Zn homeostasis and their specific contributions to pathogenesis were, until now, poorly understood. Previously, our group showed that *E. faecalis* encodes three high-affinity Mn transporters, and that while it was necessary to inactivate all three Mn transport systems (*efaCBA*, *mntH1* and *mnthH2*) to severely impair Mn uptake *in vitro*, the inactivation of only two of them (*efaCBA* and *mntH2*) was sufficient to abolish *E. faecalis* virulence (41). In this report, we showed that AdcACB and the orphan AdcAII, both predicted to mediate Zn import, are indeed critical for the ability of *E. faecalis* to grow and survive Zn-restricted conditions. Though the differences were not as striking as those seen with the Mn transport mutants (41); virulence of strains lacking the AdcACB/AdcAII system was significantly attenuated in both invertebrate and vertebrate infection models.

During characterization of the *adc* mutants, we noted that simultaneous deletion of *adcACB* and *adcAII* led to readily discernible morphological and biophysical alterations, i.e. increased cell chaining and cell-cell aggregation, which led us to wonder if the inability to maintain Zn homeostasis affected cell envelope homeostasis. Indeed, the Δ*adcA*Δ*adcAII,* Δ*adcACB* and Δ*adcACB*Δ*adcAII* strains showed heightened sensitivity to ampicillin, bacitracin and daptomycin, the latter showing a striking 16-fold lower MIC than the WT strain MIC (16 μg ml^- 1^ml, compared to 256 μg ml^-1^). While the roles played by Zn in *E. faecalis* envelope homeostasis are unknown, previous studies have associated loss of Zn transporters or Zn-dependent enzymes to surface-associated defective phenotypes. For example, in *S. pneumoniae*, deletion of *adcACB* and *adcAII* resulted in asymmetrical septa formation, abnormal cell division patterns and emergence of small aborted cells when *S. pneumoniae* was forced to grown in Zn-restricted condition (16). In the distantly-related Gram-negative pathogen *Acinetobacter baumannii*, inactivation of a Zn-dependent peptidase, ZrlA, increased cell permeability and susceptibility to the β-lactam antibiotic carbenicillin (51). Considering that tolerance to daptomycin in *E. faecalis* is, in great part, associated with structural changes at the cell envelope interface, which includes altered membrane phospholipid content and fatty acid profile that decrease membrane fluidity and the capacity of daptomycin to depolarize the cell membrane (52–54), we compared the fatty acid composition of WT and *ΔadcACBΔadcAII* strains. The total extracted lipid content of the *ΔadcACBΔadcAII* was ∼ 20% higher than the WT with the mutant displaying significant increases in palmitic acid (∼ 20% increase) and oleic acid (∼ 43% increase), the most abundant saturated and unsaturated fatty acids, respectively. While saturated fatty acids such as palmitic acid are linked with increased membrane fluidity, previous studies from another group revealed that incorporation of the unsaturated fatty acids oleic acid and linoleic acid from the environment enhanced tolerance of *E. faecalis* to daptomycin (55, 56). While not an abundant unsaturated fatty acid as oleic acid, linoleic acid concentrations were also increased (∼ 88% increase) in the *ΔadcACBΔadcAII* strain, such that concentrations of the two unsaturated fatty acids associated with daptomycin tolerance are significantly elevated in *ΔadcACBΔadcAII*. While this finding offers an explanation for the daptomycin hyper-tolerance as observed in some of the *adc* mutants, how the loss of Zn homeostasis is linked to an altered membrane fatty acid profile remains to be determined.

Similar to streptococci, enterococcal genomes do not encode the machinery to synthesize opine-like zincophores and their cognate transporters. In these Gram-positive cocci, Zn acquisition is primarily mediated by AdcABC/ZnuABC, which is under the putative control of the AdcR/ZuR repressor (22, 25, 57). However, reportedly in streptococci, a homolog of the AdcA lipoprotein, *adcAII*, is also regulated by AdcR but located far from *adcABC* and *adcR* in the chromosome (16, 26, 29). Similar to our findings showing the additive contribution of *E. faecalis adcA* and *adcAII* genes to Zn homeostasis, simultaneous inactivation of both *adcA* and *adcAII* is necessary to (nearly) abolish Zn import and, as a result, drastically impairs virulence of major human pathogens such as *S. pyogenes* and *S. pneumoniae* (16, 26, 27). Although *E. faecalis* AdcA and AdcAII share 52% identity, they do not have a similar domain organization. Of interest, biochemical and biophysical characterizations of *S. pneumoniae* AdcA and AdcAII proteins indicated that these functionally-redundant proteins employ distinct Zn acquisition mechanisms (17). AdcA has two Zn-binding domains, an amino-terminal cluster A-I domain typical of solute-binding proteins and a C-terminal domain that is structurally related to ZinT, a periplasmic Zn chaperone of Gram-negative bacteria, whereas AdcAII has only the terminal cluster A-I domain (17). Moreover, AdcAII-mediated Zn uptake *in vivo* has been shown to depend on proteins with poly-histidine (Pht) triad (HxxHxH) motifs that scavenge extracellular Zn and then transfer it to AdcAII for internalization (58). While streptococcal species have been shown to encode as many as four Pht protein homologues (e.g., *S. pneumoniae* PhtA, PhtB, PhtD and PhtE), *E. faecalis* genomes do not contain genes with HxxHxH motifs. Of note, a small open-reading frame (*OG1RF_RS12620*) (40 amino acids) coding for a putative uncharacterized protein is located upstream and separated by only 21-bp from the *adcAII* start codon. While the predicted amino acid sequence of the *OG1RF_RS12620* contains only a single histidine residue and is not enriched for other amino acids (such as cysteine and methionine) that typically coordinate Zn, it will be interesting to explore the possible role of *OG1RF_RS12620* in Zn acquisition in future studies.

Because a strain lacking the entire *adcACB* operon is phenotypically similar to the *ΔadcAΔadcAII* mutant, an important finding from this study is that both AdcA and AdcAII are capable to form functional complexes with AdcB (inner membrane permease) and AdcC (cytoplasmic ATPase). Also of interest was the distinct importance of AdcA and AdcAII in colonization and systemic dissemination observed in the two mouse models. While the attenuated virulence of *ΔadcA* and *ΔadcAII* single mutants were comparable in the peritonitis model, AdcAII was dispensable for bladder and catheter colonization in the CAUTI model. In addition, the recovery of viable bacteria from kidney and spleen of mice infected with Δ*adcAII* indicates, at first glance, suggested that the loss of AdcAII promotes bacterial dissemination to kidneys and spleen. While speculative at this point, we believe that these unexpected phenotypes are due to differences in expression levels of *adcA* and *adcAII* in the bladder environment. We propose that *adcA* responds more strongly than *adcAII* to environmental cues encountered in the bladder, such that *adcAII* becomes dispensable for *E. faecalis* proliferation in urine. To investigate this possibility, studies to compare the transcriptional patterns of *adcACB* and Δ*adcAII* in the bladder, peritoneal cavity and bloodstream environments will be performed in the near future. Alternatively, the large and constant fluctuations in the bladder environment in solute concentrations and of other important biophysical and biochemical parameters caused by intermittent cycles of urination can somehow compromise Zn-binding capacity of AdcAII or ability to interact with the AdcBC partner proteins.

In summary, our findings reveal that the AdcACB/AdcAII system is a bona fide Zn acquisition system of *E. faecalis* contributing additively in the maintenance of Zn homeostasis. More importantly, this report demonstrates that the AdcACB/AdcAII system mediates *E. faecalis* virulence. Therefore, both the substrate-binding and surface-associated AdcA and AdcAII proteins can be viewed as suitable targets for the development of antimicrobial therapies to treat or prevent enterococcal infections.

## MATERIAL & METHODS

### Bacterial strains and growth conditions

The strains and plasmids used in this study are listed in Table 2. Unless stated otherwise, bacteria were grown in brain heart infusion (BHI broth (BD Difco^TM^, for *E. faecalis*) and Luria-bertani (BD Difco^TM^, for *E. coli*) at 37°C under static conditions. Strains harboring the pGCP123 plasmid (59) were grown in the presence of kanamycin (300 µg ml^-1^ for *E. coli*, 500 µg ml^-1^ for *E. faecalis*). For growth kinetics assays, overnight cultures were normalized to an OD_600_ of 0.25 and inoculated into fresh BHI at a 1:50 ratio, with the OD_600_ monitored over time in an automated growth reader (Bioscreen c, Oy Growth Curves AB). Purified wild-type (Cp) Mn-deficient calprotectin (Cp ^ΔMn-tail^) calprotectin were gifts from Dr. Walter Chazin (Vanderbilt University, USA)(10). Experiments using calprotectin were performed in BHI supplemented with 20% (v/v) CP buffer (40 mM NaCl, 0.5 mM β-mercapethanol, 1.2 mM CaCl_2_, 8 mM Tris-HCl, pH 7.5). Normalization of starter cultures for most experiments included centrifugation or overnight cultures at 4000 rpm for 10 mins to remove spent media, two cell pellet washes in PBS, and adjustment of cell density to an OD_600_ of 0.25 (∼ 1 x 10^8^ CFU ml^-1^) in PBS. For *E. faecalis* CFU determination from *ex vivo* and *in vivo* studies, serially-diluted aliquots were plated on BHI agar with 200 µg ml^-1^ rifampicin and 10 µg ml^-1^ fusidic acid. TPEN (N,N,N′,N′-tetrakis(2-pyridinylmethyl)-1,2-ethanediamine), FeSO_4_, MnSO_4_, ZnSO_4_, lysozyme, ampicillin, bacitracin, daptomycin, fusidic acid, kanamycin, rifampicin and vancomycin were purchased from Sigma Aldrich.

**Table 2.**
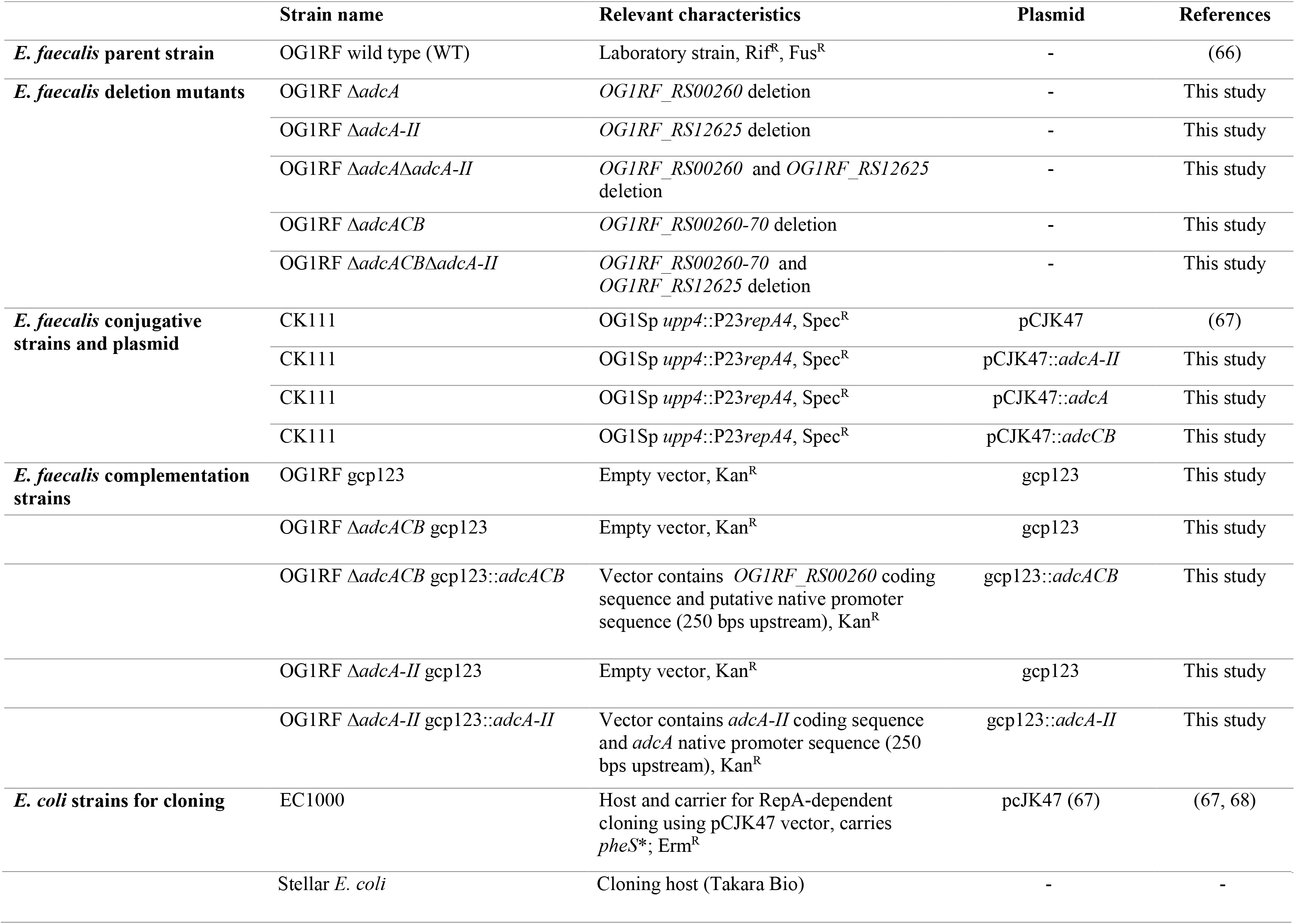
Strains and plasmids used in this study.

|  | Strain name | Relevant characteristics | Plasmid | References |
| --- | --- | --- | --- | --- |
| *E. faecalis* parent strain | OG1RF wild type (WT) | Laboratory strain, Rif<sup>R</sup>, Fus<sup>R</sup> | - | (66) |
| *E. faecalis* deletion mutants | OG1RF Δ*adcA* | *OG1RF_RS00260* deletion | - | This study |
| | OG1RF Δ*adcA-II* | *OG1RF_RS12625* deletion | - | This study |
| | OG1RF Δ*adcA*Δ*adcA-II* | *OG1RF_RS00260* and *OG1RF_RS12625* deletion | - | This study |
| | OG1RF Δ*adcACB* | *OG1RF_RS00260-70* deletion | - | This study |
| | OG1RF Δ*adcACB*Δ*adcA-II* | *OG1RF_RS00260-70* and *OG1RF_RS12625* deletion | - | This study |
| *E. faecalis* conjugative strains and plasmid | CK111 | OG1Sp *upp4*::P23*repA4*, Spec<sup>R</sup> | pCJK47 | (67) |
|  | CK111 | OG1Sp *upp4*::P23*repA4*, Spec<sup>R</sup> | pCJK47::*adcA-II* | This study |
|  | CK111 | OG1Sp *upp4*::P23*repA4*, Spec<sup>R</sup> | pCJK47::*adcA* | This study |
|  | CK111 | OG1Sp *upp4*::P23*repA4*, Spec<sup>R</sup> | pCJK47::*adcCB* | This study |
| *E. faecalis* complementation strains | OG1RF gcp123 | Empty vector, Kan<sup>R</sup> | gcp123 | This study |
| | OG1RF Δ*adcACB* gcp123 | Empty vector, Kan<sup>R</sup> | gcp123 | This study |
| | OG1RF Δ*adcACB* gcp123::*adcACB* | Vector contains *OG1RF_RS00260* coding sequence and putative native promoter sequence (250 bps upstream), Kan<sup>R</sup> | gcp123::*adcACB* | This study |
| | OG1RF Δ*adcA-II* gcp123 | Empty vector, Kan<sup>R</sup> | gcp123 | This study |
| | OG1RF Δ*adcA-II* gcp123::*adcA-II* | Vector contains *adcA-II* coding sequence and *adcA* native promoter sequence (250 bps upstream), Kan<sup>R</sup> | gcp123::*adcA-II* | This study |
| *E. coli* strains for cloning | EC1000 | Host and carrier for RepA-dependent cloning using pCJK47 vector, carries *pheS* *; Erm<sup>R</sup> | pCJK47 (67) | (67, 68) |

|  |  |  |  |
| --- | --- | --- | --- |
| Stellar *E. coli* | Cloning host (Takara Bio) | - | - |

### General cloning techniques

The nucleotide sequences of *adcACB* and *adcAII* were obtained from the *E. faecalis* OG1RF genome available at BioCyc (60). Wizard genomic DNA purification kit (Promega, USA) was used for isolation of bacterial genomic DNA (gDNA), and Monarch plasmid miniprep kit (New England BioLabs) used for plasmid purification. The Monarch DNA gel extraction kit (New England BioLabs) was used to isolate PCR products. The In-Fusion HD cloning kit (TaKaRa Bio) was used for fast, directional cloning of DNA fragments into expression vectors. Colony PCR was performed using PCR 2x Master Mix (Promega) with primers listed in Table 3.

**Table 3.**
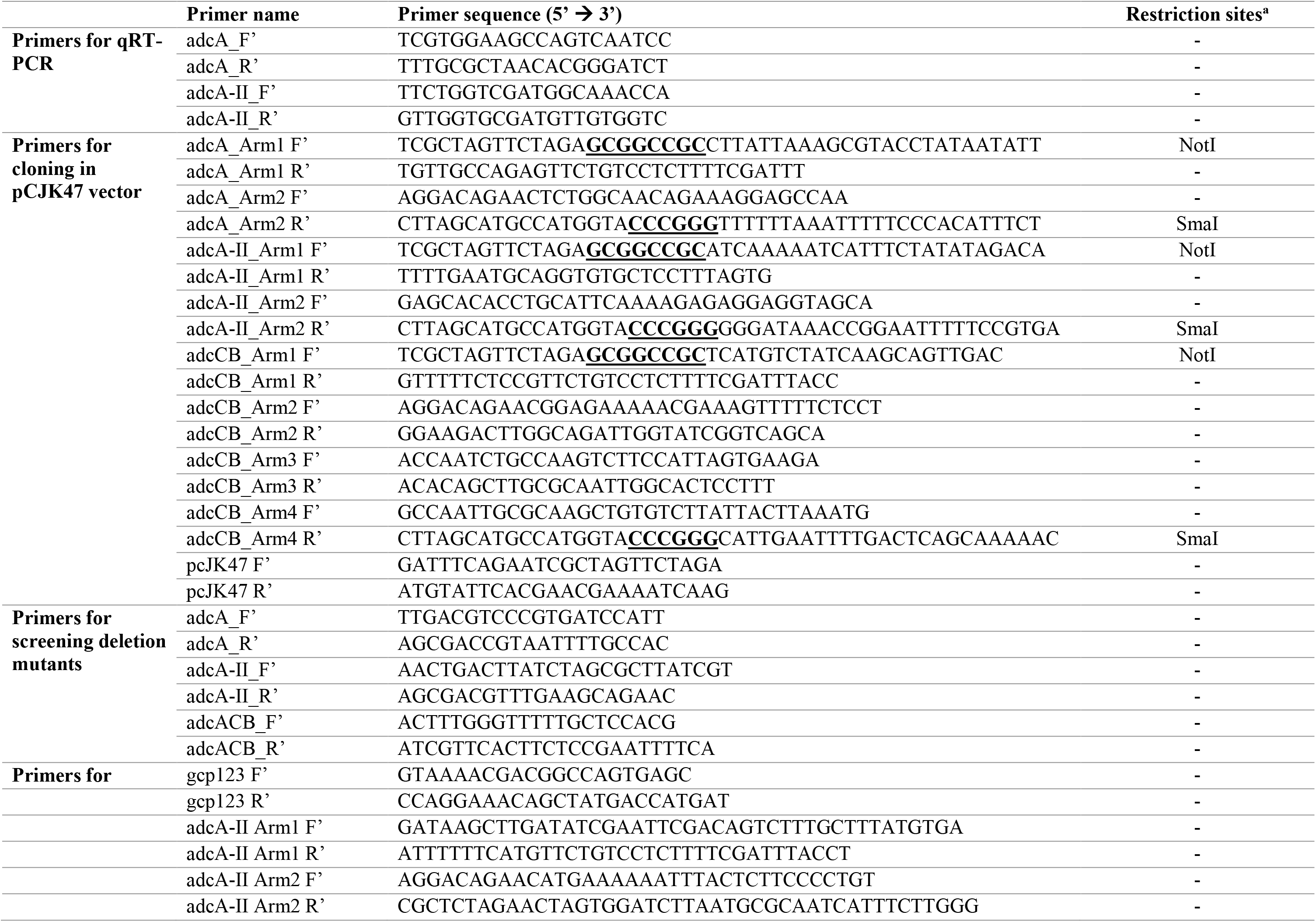
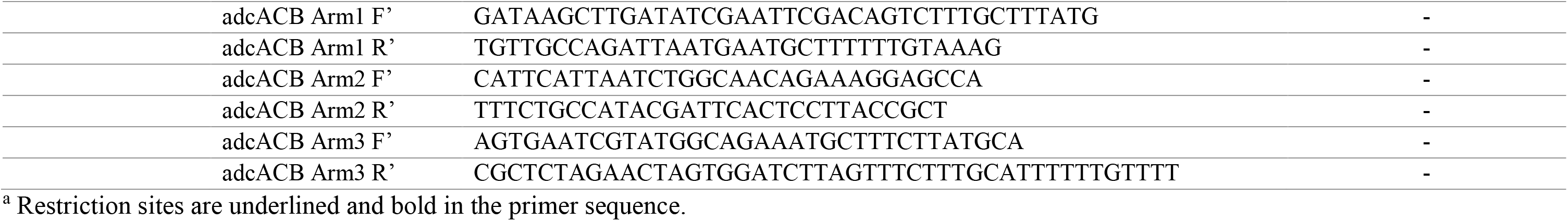
Primers used in this study.

### Construction of deletion and genetically complemented strains

Deletion of *adcA*, *adcAII* or the entire *adcACB* operon was carried out using the pCJK47 markerless genetic exchange system (38). Briefly, upstream and downstream nucleotide sequences of *adcA, adcCB* and *adcAII* were amplified using the primers listed in Table 3 to generate amplicon templates. Introduction of amplicons into the pcJK47 vector, followed by conjugation into *E*. *faecalis* OG1RF and isolation of deletion mutants were carried out as previously described (38). The Δ*adcA*Δ*adcAII* and Δ*adcACB*Δ*adcAII* double mutants were obtained by conjugating the pCJK-*adcAII* plasmid with the Δ*adcA* and Δ*adcACB* mutants, respectively. Because the *adcACB* operon is ∼ 3kb, we first isolated the Δ*adcA* mutant and then performed conjugation using the pCJK-*adcBC* vector to generate the Δ*adcACB* strain. All gene deletions were confirmed by PCR sequencing of the insertion site and flanking sequences. For genetic complementation, full-length *adcACB* and *adcAII* genes were amplified by PCR using the primers listed in Table 3, and cloned onto the pGCP123 plasmid (59) digested with BamHI and EcoRI restriction enzymes. The full-length *adcACB* and *adcAII* nucleotide sequences were incorporated into linearized pGCP123 plasmid using In-Fusion HD cloning kit and transformed into stellar *E. coli*. Plasmids were propagated in *E. coli* and electroporated into *E. faecalis* OG1RF and mutant derivatives as described previously (59).

### Inductively coupled plasma – optical emission spectrometry (ICP-OES)

Trace metal concentration in BHI media and in bacteria was determined as previously described (41). For quantification of metals in BHI, 9 ml of BHI was digested with 1 ml of trace-metal grade 35% nitric acid (HNO_3_) prior to analysis. For quantification of cellular metal content, overnight cultures were washed twice with PBS and subsequently used to inoculate into fresh BHI or BHI supplemented with 7.5 µM TPEN at a ratio of 1:40. After reaching an OD_600_ of 0.5, cells were harvested by centrifugation, washed twice with PBS containing 0.5 mM EDTA and aliquots collected for metal analysis. Cell pellets were resuspended in 1 ml trace-metal grade 35% nitric acid (HNO_3_) and digested at 90°C for 1 hour in a high-density polyethylene scintillation vial (Fisher Scientific). Digested bacterial cells were diluted at a ratio of 1:10 in reagent-grade water prior to analysis. Metal composition was quantified using a 5300DV ICP Atomic Emission Spectrometer (Perkin Elmer) and concentrations were determined by comparison with a standard curve at the University of Florida Institute of Food and Agricultural Sciences Analytical Services Laboratories. Metal concentrations were normalized to total protein content determined using the bicinchoninic acid (BCA) assay kit (Pierce^TM^).

### MIC determinations

Strains were grown overnight in BHI, normalized to an OD_600_ of 0.25, and diluted at a ratio of 1:1000. The diluted cultures were inoculated at a ratio of 1:20 into fresh BHI containing increasing concentrations of antibiotics (ampicillin, bacitracin, daptomycin, and vancomycin). The absorbance at OD_600_ after incubation at 37 °C for 24 hours was measured in a Synergy H1 microplate reader (Molecular Devices, USA).

### Growth and survival in serum and urine

Strains were grown overnight in BHI, normalized to an OD_600_ of 0.5, and inoculated at a ratio of 1:1000 into pooled human serum or pooled human urine (both purchased from Lee Biosolutions). At selected time points, aliquots were serially diluted in PBS and dilutions plated on BHI agar supplemented with 10 µg ml^-1^ fusidic acid and 200 µg ml^-1^ rifampicin for CFU determination.

### Biofilm assay

Overnight cultures grown in BHI were normalized to OD_600_ of 0.5, and further diluted 1:25 in fresh BHI supplemented with 10 mM glucose and added to the wells of 96-well polystyrene plates (Grenier CellSTAR) that were incubated under static conditions at 37°C for 24 hours. After incubation, media with planktonic cells were discarded, and the wells gently washed with PBS twice. Adherent cells were stained with 0.1% crystal violet for 25 min, and the bound dye extracted from the wells using a 33% acetic acid solution. Absorbance was measured at an OD_595_ using the Synergy H1 microplate reader.

### Quantitative real-time PCR

Strains were grown overnight in BHI, normalized to an OD_600_ of 0.5, inoculated at a ratio of 1:20 into fresh BHI and incubated at 37°C for 1 hr. Post incubation, cells were harvested via centrifugation at 4000 rpm for 10 mins, washed once with PBS and incubated with lysozyme (20 mg ml^-1^) for 30 min at 37°C. Post lysozyme treatment, cells were harvested by centrifugation and total RNA extracted in a Purifier filtered PCR enclosure using the PureLink RNA minikit (Invitrogen), according to the manufacturer’s instructions. RNA purification and removal of DNA were performed using a Turbo DNA-free kit (Thermo Fisher). Synthesis of cDNA was performed using a High-capacity cDNA Reverse Transcription Kit (Applied Biosystems). Quantitative real-time PCR (qRT-PCR) was performed using iTaq Universal SYBR supermix (BioRad) with the primers listed in Table 2. For quantification of transcript numbers, *E. faecalis* OG1RF gDNA was used as template to generate standard curves.

### Galleria mellonella infection

Larvae of *G*. *mellonella* were used as model to assess virulence of *E. faecalis* as described previously (41, 61), with minor modifications. Briefly, groups of 15 larvae (200–300 mg in weight) were injected with 5 μl of normalized overnight bacterial inoculum containing ∼ 5 x 10^5^ CFU. Larvae injected with heat-inactivated *E*. *faecalis* (30 mins at 100°C) or PBS were used as negative and vehicle controls, respectively. Post-injection, larvae were kept at 37°C and survival recorded at selected intervals for up to 72 hours.

### Catheter-associated peritonitis mouse model

The methods for the catheter-associated peritonitis model have been described previously (62), hence only minor modifications are described here. Female C57BL/6J 8-weeks old mice (Jackson Laboratories) were anesthetized and a sterile 1-cm long catheter segment (Qosina) inserted into the peritoneum using an 18-gauge BD Spinal needle/stylette (Becton, Dickinson and Company,). Three hours post-catheter implantation, mice were injected via i.p. with normalized overnight bacterial inoculum containing ∼ 5×10^8^ CFU of bacteria suspended in PBS. Forty eight hours post-infection, animals were euthanized and bacterial burden in peritoneal wash, catheter and spleen determined by plating serial dilutions of homogenates on BHI plates selective (200 µg ml^-1^ rifampicin and 10 µg ml^-1^ fusidic acid) for *E. faecalis* OG1RF and derivatives. This procedure was approved and performed in compliance with the University of Florida Institutional Animal Care and Use Committee (IACUC).

### Mouse catheter-associated urinary tract infection (CAUTI) model

The methods for the CAUTI mouse model have also been described (63). Female C57BL/6Ncr 6-weeks old mice (Charles River Laboratories) were anesthetized and subjected to transurethral implantation of a 5-mm length platinum-cured silicone catheter. Immediately after catheter implantation, mice were infected with **∼** 1×10^7^ CFU of bacteria prepared in PBS. Twenty four hours post-infection, mice were euthanized and catheter and organs collected for bacterial enumeration by plating catheter and organ homogenates on BHI plates selective for *E. faecalis* OG1RF and derivatives. This procedure is in compliance with the University of Notre Dame Institutional Animal Care and Use Committee (protocol #18-08-4792MD).

### Bright field microscopy

Overnight cultures grown in BHI were normalized to OD_600_ 0.5 (∼ 5×10^8^ CFU/ml) and washed once with 1 ml of PBS. Aliquot of 5 µl of bacterial inoculum was placed on microscope glass sides that were washed (once in filtered 70% ethanol followed by milliQ water) and dried, and left to dry at room temperature. Samples were covered with 5 µl of mounting media (Vectashield, USA) and covered with a glass coverslip. Bright field microscopy was performed using Leica DM2500 LED optical microscope fitted with a 100X/1.3 oil objective lens and Leica acquisition software. Images acquired were further processed using FIJI software (64).

### Gas chromatography fatty acid methyl ester (GC-FAME) analysis

Normalized overnight bacterial cultures (OD_600_ 0.25, ∼ 10^8^ CFU ml^-1^) were inoculated at a ratio of 1:20 into 1 L of fresh BHI and incubated for 4 hours at 37°C. After incubation, cells were harvested by centrifugation, washed twice in PBS, and the cell pellets frozen at −80°C prior to use. Frozen samples were processed for GC-FAME analysis at Analytical Toxicology Core Laboratory, Center for Environmental and Human Toxicology, University of Florida. A modified direct derivatization procedure (65) involved addition of 14% BF_3_ in methanol to samples, followed by incubation in a 100° C sand bath and back-extraction with hexane (Fisher Scientific). Sample extracts were concentrated to 0.3 ml under a gentle stream of N_2_ at 35°C. US-108N mixed deuterated PAH standard (Agilent) was added to the samples as an internal standard prior to analysis. Authentic FAME standards were prepared as a set of dilutions 0.01 – 5.0 µg ml^-1^ (GLC-461 standard mix, Nu-Chek Prep) spiked with US-108N ISTD for FAME quantitation. Sample testing was carried out on an Agilent 7890B gas chromatograph equipped with a DB-FATWAX UI column (30 m x 0.25 mm ID x 0.25 µm film thickness, Agilent), and coupled to an Agilent 7000C mass spectrometer. The sample inlet was 280° C, with constant He flow at 1.2 mL/min. The oven temperature program was as follows: initial temperature 40° C held 4 min, Ramp 1 increase 10° C/min to 180° C and held 5 min, Ramp 2 increase 5° C/min to 250° C and held 3 min. Sample components were then passed through a 280° C transfer line into the mass spectrometer. The MS source temperature was 230° C and quad 1 at 150° C. The mode of data acquisition was in both full-scan and in selected ion monitoring (SIM); SIM mode was used for low-concentration lipid species, and for distinguishing co-eluting C22:6n3 and C24:1n9. Quantification of fatty acid species was normalized against total weight of cell pellet processed.

### Statistical analysis

Data were analysed using GraphPad Prism 9.0 software (GraphPad Software, San Diego, CA, USA). Data from multiple experiments were pooled, and appropriate statistical tests were applied, as indicated in the figure legends. An adjusted value of *p* ≤ 0.05 was considered statistically significant.

## ACKNOWLEDGEMENT

This study was supported by NIH/NIAID R21 AI135158 to J.A.L. and by institutional funds from the University of Notre Dame and NIH/NIDDK R01 DK128805 to J.J.M. and A.L.F.M. D.N.B. was supported by NIH/NIDCR T90 DE021990.

## DISCLOSURE OF POTENTIAL CONFLICTS OF INTEREST

The authors declare that they have no competing interests.

## DATA AVAILABILITY STATEMENT

The authors confirm that the data supporting the findings of this study are available within the article and/or its supplementary materials. Vectors and strains created from this study will be available from the corresponding author, upon reasonable request.

## SUPPLEMENTARY DATA

**Fig S1.**
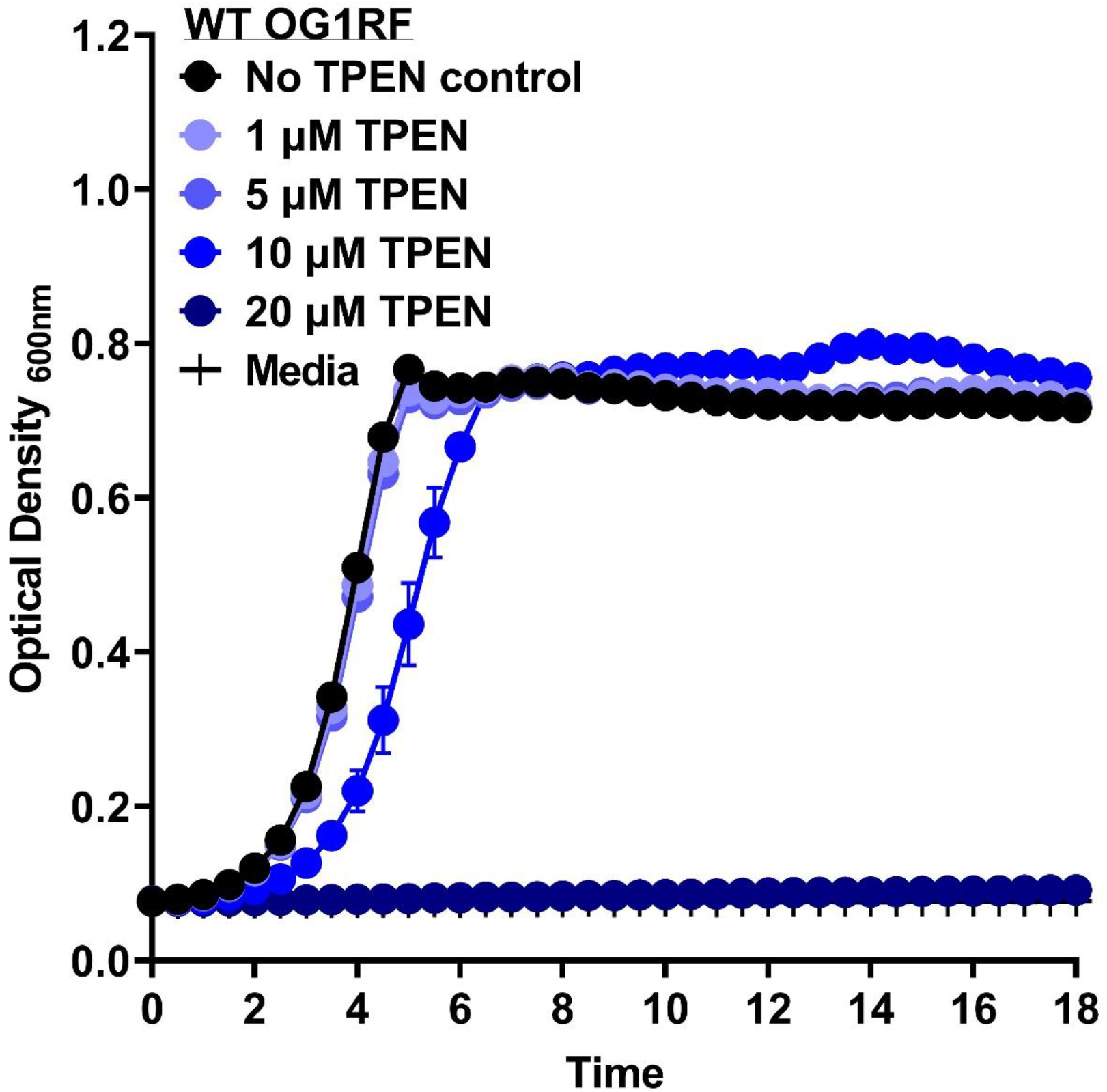
Growth characteristics of *E. faecalis* in the presence of TPEN. Growth curves of *E. faecalis* OG1RF (WT) in BHI supplemented with increasing concentrations of TPEN. Data points represent the average and error bar represent the standard error of margin (SEM) of nine biological replicates. Statistical analysis was performed using simple linear regression of exponential growth phase, and slope of each growth kinetics was compared to the parent strain grown without TPEN.

**Fig S2.**
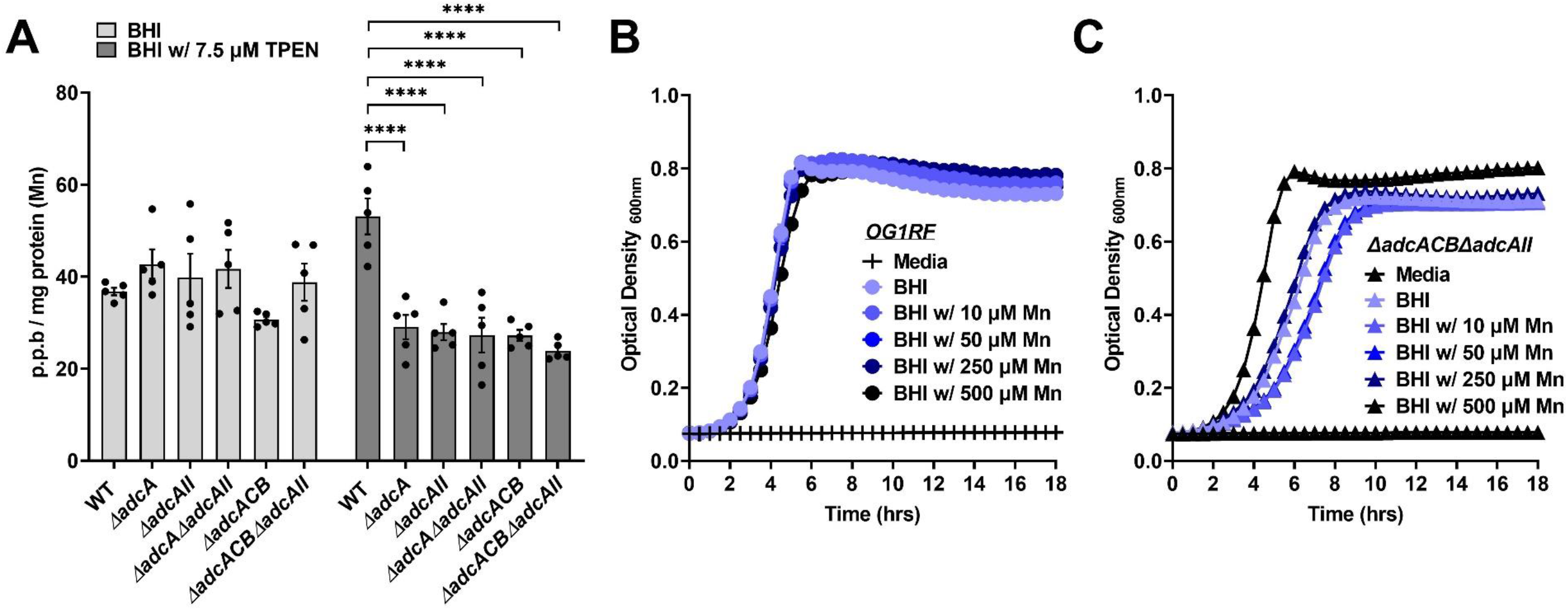
Phenotypic characterization of intracellular Mn in *E. faecalis* Zn-deficient mutants. (A) ICP-OES quantification of intracellular Mn of mid-log grown *E. faecalis* OG1RF (WT) and derivatives grown in BHI or BHI supplemented with 7.5 µM TPEN. Data points represent five biological replicates. Statistical analysis was performed using two-way ANOVA with Dunnett’s multiple comparison test. Error bar represent the standard error of margin (SEM). ** *p* ≤ 0.01, *** *p* ≤ 0.001 and **** *p* ≤ 0.0001. (B-C) Growth curves of *E. faecalis* OG1R (B) and *ΔadcACBΔadcAII* (C) in BHI supplemented with increasing Mn concentrations. Data points represent six biological replicates. Statistical analysis was performed using simple linear regression of exponential growth phase, and slope of each growth kinetic was compared to the reference strain grown without Mn supplementation.

## Notes

### Competing Interest Statement

The authors have declared no competing interest.

## REFERENCES

1. Andreini C, Banci L, Bertini I, Rosato A. 2006. Counting the zinc-proteins encoded in the human genome. J Proteome Res 5:196–201.

2. Hood MI, Skaar EP. 2012. Nutritional immunity: transition metals at the pathogen–host interface. Nature Reviews Microbiology 10:525–537.

3. Gammoh NZ, Rink L. 2017. Zinc in Infection and Inflammation. Nutrients 9.

4. Andreini C, Banci L, Bertini I, Rosato A. 2006. Zinc through the three domains of life. J Proteome Res 5:3173–8.

5. Kehl-Fie TE, Skaar EP. 2010. Nutritional immunity beyond iron: a role for manganese and zinc. Curr Opin Chem Biol 14:218–24.

6. Cassat JE, Skaar EP. 2013. Iron in infection and immunity. Cell Host Microbe 13:509–519.

7. Juttukonda LJ, Skaar EP. 2015. Manganese homeostasis and utilization in pathogenic bacteria. Mol Microbiol doi:10.1111/mmi.13034.

8. Palmer LD, Skaar EP. 2016. Transition Metals and Virulence in Bacteria. Annu Rev Genet 50:67–91.

9. Zygiel EM, Nolan EM. 2018. Transition Metal Sequestration by the Host-Defense Protein Calprotectin. Annu Rev Biochem 87:621–643.

10. Damo SM, Kehl-Fie TE, Sugitani N, Holt ME, Rathi S, Murphy WJ, Zhang Y, Betz C, Hench L, Fritz G, Skaar EP, Chazin WJ. 2013. Molecular basis for manganese sequestration by calprotectin and roles in the innate immune response to invading bacterial pathogens. Proc Natl Acad Sci U S A 110:3841–6.

11. Becker KW, Skaar EP. 2014. Metal limitation and toxicity at the interface between host and pathogen. FEMS Microbiol Rev 38:1235–49.

12. Makthal N, Nguyen K, Do H, Gavagan M, Chandrangsu P, Helmann JD, Olsen RJ, Kumaraswami M. 2017. A Critical Role of Zinc Importer AdcABC in Group A Streptococcus-Host Interactions During Infection and Its Implications for Vaccine Development. EBioMedicine 21:131–141.

13. Mastropasqua MC, D’Orazio M, Cerasi M, Pacello F, Gismondi A, Canini A, Canuti L, Consalvo A, Ciavardelli D, Chirullo B, Pasquali P, Battistoni A. 2017. Growth of Pseudomonas aeruginosa in zinc poor environments is promoted by a nicotianamine-related metallophore. Mol Microbiol 106:543–561.

14. Lhospice S, Gomez NO, Ouerdane L, Brutesco C, Ghssein G, Hajjar C, Liratni A, Wang S, Richaud P, Bleves S, Ball G, Borezée-Durant E, Lobinski R, Pignol D, Arnoux P, Voulhoux R. 2017. Pseudomonas aeruginosa zinc uptake in chelating environment is primarily mediated by the metallophore pseudopaline. Scientific reports 7:17132–17132.

15. Paik S, Brown A, Munro CL, Cornelissen CN, Kitten T. 2003. The sloABCR operon of Streptococcus mutans encodes an Mn and Fe transport system required for endocarditis virulence and its Mn-dependent repressor. J Bacteriol 185:5967–75.

16. Bayle L, Chimalapati S, Schoehn G, Brown J, Vernet T, Durmort C. 2011. Zinc uptake by Streptococcus pneumoniae depends on both AdcA and AdcAII and is essential for normal bacterial morphology and virulence. Mol Microbiol 82:904–16.

17. Plumptre CD, Eijkelkamp BA, Morey JR, Behr F, Counago RM, Ogunniyi AD, Kobe B, O’Mara ML, Paton JC, McDevitt CA. 2014. AdcA and AdcAII employ distinct zinc acquisition mechanisms and contribute additively to zinc homeostasis in Streptococcus pneumoniae. Mol Microbiol 91:834–51.

18. Capdevila DA, Wang J, Giedroc DP. 2016. Bacterial Strategies to Maintain Zinc Metallostasis at the Host-Pathogen Interface. J Biol Chem 291:20858–20868.

19. Frassinetti S, Bronzetti G, Caltavuturo L, Cini M, Croce CD. 2006. The role of zinc in life: a review. J Environ Pathol Toxicol Oncol 25:597–610.

20. Lonergan ZR, Skaar EP. 2019. Nutrient Zinc at the Host-Pathogen Interface. Trends Biochem Sci 44:1041–1056.

21. Crawford A, Wilson D. 2015. Essential metals at the host-pathogen interface: nutritional immunity and micronutrient assimilation by human fungal pathogens. FEMS Yeast Res 15.

22. Blindauer CA. 2015. Advances in the molecular understanding of biological zinc transport. Chem Commun (Camb) 51:4544–63.

23. Morey JR, Kehl-Fie TE. 2020. Bioinformatic Mapping of Opine-Like Zincophore Biosynthesis in Bacteria. mSystems. 5(4):doi:10.1128/msystems.00554-20.

24. Grim KP, San Francisco B, Radin JN, Brazel EB, Kelliher JL, Párraga Solórzano PK, Kim PC, McDevitt CA, Kehl-Fie TE. 2017. The Metallophore Staphylopine Enables Staphylococcus aureus To Compete with the Host for Zinc and Overcome Nutritional Immunity. mBio 8.

25. Makthal N, Kumaraswami M. 2017. Zinc’ing it out: zinc homeostasis mechanisms and their impact on the pathogenesis of human pathogen group A streptococcus. Metallomics 9:1693–1702.

26. Makthal N, Do H, Wendel BM, Olsen RJ, Helmann JD, Musser JM, Kumaraswami M. 2020. Group A Streptococcus AdcR Regulon Participates in Bacterial Defense against Host-Mediated Zinc Sequestration and Contributes to Virulence. Infect Immun 88.

27. Ong CY, Berking O, Walker MJ, McEwan AG. 2018. New Insights into the Role of Zinc Acquisition and Zinc Tolerance in Group A Streptococcal Infection. Infect Immun 86.

28. Burcham LR, Le Breton Y, Radin JN, Spencer BL, Deng L, Hiron A, Ransom MR, Mendonça JDC, Belew AT, El-Sayed NM, McIver KS, Kehl-Fie TE, Doran KS. 2020. Identification of Zinc-Dependent Mechanisms Used by Group B Streptococcus To Overcome Calprotectin-Mediated Stress. mBio 11.

29. Ganguly T, Peterson AM, Kajfasz JK, Abranches J, Lemos JA. 2021. Zinc Import Mediated by AdcABC is Critical for Colonization of the Dental Biofilm by Streptococcus mutans in an Animal Model. bioRxiv doi:10.1101/2021.01.22.427828:2021.01.22.427828.

30. Fiore E, Van Tyne D, Gilmore MS. 2019. Pathogenicity of Enterococci. Microbiology spectrum 7:10.1128/microbiolspec.GPP3-0053-2018.

31. Ch’ng J-H, Chong KKL, Lam LN, Wong JJ, Kline KA. 2019. Biofilm-associated infection by enterococci. Nature Reviews Microbiology 17:82–94.

32. Vu J, Carvalho J. 2011. Enterococcus: review of its physiology, pathogenesis, diseases and the challenges it poses for clinical microbiology. Frontiers in Biology 6:357.

33. Gilmore MS CD, Ike Y, et al (ed). 2014. Enterococci: From Commensals to Leading Causes of Drug Resistant Infection. Boston: Massachusetts Eye and Ear Infirmary, Accessed

34. Miller WR, Munita JM, Arias CA. 2014. Mechanisms of antibiotic resistance in enterococci. Expert review of anti-infective therapy 12:1221–1236.

35. Gaca AO, Lemos JA. 2019. Adaptation to Adversity: the Intermingling of Stress Tolerance and Pathogenesis in Enterococci. Microbiology and Molecular Biology Reviews 83:e00008–19.

36. Coelho Abrantes M, Lopes MdF, Kok J. 2011. Impact of Manganese, Copper and Zinc Ions on the Transcriptome of the Nosocomial Pathogen Enterococcus faecalis V583. PLOS ONE 6:e26519.

37. Latorre M, Low M, Gárate E, Reyes-Jara A, Murray BE, Cambiazo V, González M. 2015. Interplay between copper and zinc homeostasis through the transcriptional regulator Zur in Enterococcus faecalis. Metallomics 7:1137–45.

38. Kristich CJ, Chandler JR, Dunny GM. 2007. Development of a host-genotype-independent counterselectable marker and a high-frequency conjugative delivery system and their use in genetic analysis of Enterococcus faecalis. Plasmid 57:131–44.

39. 39. Kajfasz JK, Katrak C, Ganguly T, Vargas J, Wright L, Peters ZT, Spatafora GA, Abranches J, Lemos JA. 2020. Manganese Uptake, Mediated by SloABC and MntH, Is Essential for the Fitness of Streptococcus mutans. mSphere 5.

40. Zhang F, Ma XL, Wang YX, He CC, Tian K, Wang HG, An D, Heng B, Xie LH, Liu YQ. 2017. TPEN, a Specific Zn(2+) Chelator, Inhibits Sodium Dithionite and Glucose Deprivation (SDGD)-Induced Neuronal Death by Modulating Apoptosis, Glutamate Signaling, and Voltage-Gated K(+) and Na(+) Channels. Cell Mol Neurobiol 37:235–250.

41. Colomer-Winter C, Flores-Mireles AL, Baker SP, Frank KL, Lynch AJL, Hultgren SJ, Kitten T, Lemos JA. 2018. Manganese acquisition is essential for virulence of Enterococcus faecalis. PLoS pathogens 14:e1007102–e1007102.

42. Roohani N, Hurrell R, Kelishadi R, Schulin R. 2013. Zinc and its importance for human health: An integrative review. Journal of research in medical sciences : the official journal of Isfahan University of Medical Sciences 18:144–157.

43. King JC, Shames DM, Woodhouse LR. 2000. Zinc homeostasis in humans. J Nutr 130:1360s-6s.

44. Maynar M, Muñoz D, Alves J, Barrientos G, Grijota FJ, Robles MC, Llerena F. 2018. Influence of an Acute Exercise Until Exhaustion on Serum and Urinary Concentrations of Molybdenum, Selenium, and Zinc in Athletes. Biological Trace Element Research 186:361–369.

45. Chan S, Gerson B, Subramaniam S. 1998. The role of copper, molybdenum, selenium, and zinc in nutrition and health. Clin Lab Med 18:673–85.

46. Cordova A, Alvarez-Mon M. 1995. Behaviour of zinc in physical exercise: a special reference to immunity and fatigue. Neurosci Biobehav Rev 19:439–45.

47. Sandstead HH. 2015. Chapter 61 - Zinc, p 1369–1385. In Nordberg GF, Fowler BA, Nordberg M (ed), Handbook on the Toxicology of Metals (Fourth Edition) doi:https://doi.org/10.1016/B978-0-444-59453-2.00061-5. Academic Press, San Diego.

48. Brewer W, Harrison J, Saito HE, Fozo EM. 2020. Induction of Daptomycin Tolerance in Enterococcus faecalis by Fatty Acid Combinations. Appl Environ Microbiol 86.

49. Mishra NN, Bayer AS, Tran TT, Shamoo Y, Mileykovskaya E, Dowhan W, Guan Z, Arias CA. 2012. Daptomycin Resistance in Enterococci Is Associated with Distinct Alterations of Cell Membrane Phospholipid Content. PLOS ONE 7:e43958.

50. Woodall BM, Harp JR, Brewer WT, Tague ED, Campagna SR, Fozo EM. 2021. Enterococcus faecalis Readily Adapts Membrane Phospholipid Composition to Environmental and Genetic Perturbation. Frontiers in Microbiology 12.

51. Lonergan ZR, Nairn BL, Wang J, Hsu Y-P, Hesse LE, Beavers WN, Chazin WJ, Trinidad JC, VanNieuwenhze MS, Giedroc DP, Skaar EP. 2019. An Acinetobacter baumannii, Zinc-Regulated Peptidase Maintains Cell Wall Integrity during Immune-Mediated Nutrient Sequestration. Cell Reports 26:2009–2018.e6.

52. Tran TT, Panesso D, Mishra NN, Mileykovskaya E, Guan Z, Munita JM, Reyes J, Diaz L, Weinstock GM, Murray BE, Shamoo Y, Dowhan W, Bayer AS, Arias CA, Projan SJ. 2013. Daptomycin-Resistant Enterococcus faecalis Diverts the Antibiotic Molecule from the Division Septum and Remodels Cell Membrane Phospholipids. mBio 4:e00281–13.

53. Ledger EVK, Pader V, Edwards AM. 2017. Enterococcus faecalis and pathogenic streptococci inactivate daptomycin by releasing phospholipids. Microbiology (Reading) 163:1502–1508.

54. Saito HE, Harp JR, Fozo EM. 2014. Incorporation of exogenous fatty acids protects Enterococcus faecalis from membrane-damaging agents. Appl Environ Microbiol 80:6527–38.

55. Saito HE, Harp JR, Fozo EM. 2017. Enterococcus faecalis Responds to Individual Exogenous Fatty Acids Independently of Their Degree of Saturation or Chain Length. Applied and environmental microbiology 84:e01633–17.

56. Harp JR, Saito HE, Bourdon AK, Reyes J, Arias CA, Campagna SR, Fozo EM, Vieille C. 2016. Exogenous Fatty Acids Protect Enterococcus faecalis from Daptomycin-Induced Membrane Stress Independently of the Response Regulator LiaR. Applied and Environmental Microbiology 82:4410–4420.

57. Mikhaylina A, Ksibe AZ, Scanlan DJ, Blindauer CA. 2018. Bacterial zinc uptake regulator proteins and their regulons. Biochemical Society Transactions 46:983–1001.

58. Bersch B, Bougault C, Roux L, Favier A, Vernet T, Durmort C. 2013. New insights into histidine triad proteins: solution structure of a Streptococcus pneumoniae PhtD domain and zinc transfer to AdcAII. PLoS One 8:e81168.

59. Nielsen HV, Guiton PS, Kline KA, Port GC, Pinkner JS, Neiers F, Normark S, Henriques-Normark B, Caparon MG, Hultgren SJ. 2012. The metal ion-dependent adhesion site motif of the Enterococcus faecalis EbpA pilin mediates pilus function in catheter-associated urinary tract infection. mBio 3:e00177–12.

60. Karp PD, Billington R, Caspi R, Fulcher CA, Latendresse M, Kothari A, Keseler IM, Krummenacker M, Midford PE, Ong Q, Ong WK, Paley SM, Subhraveti P. 2019. The BioCyc collection of microbial genomes and metabolic pathways. Brief Bioinform 20:1085–1093.

61. Gaca AO, Abranches J, Kajfasz JK, Lemos JA. 2012. Global transcriptional analysis of the stringent response in Enterococcus faecalis. Microbiology 158:1994–2004.

62. Kajfasz JK, Mendoza JE, Gaca AO, Miller JH, Koselny KA, Giambiagi-Demarval M, Wellington M, Abranches J, Lemos JA. 2012. The Spx regulator modulates stress responses and virulence in Enterococcus faecalis. Infection and immunity 80:2265–2275.

63. Flores-Mireles AL, Pinkner JS, Caparon MG, Hultgren SJ. 2014. EbpA vaccine antibodies block binding of Enterococcus faecalis to fibrinogen to prevent catheter-associated bladder infection in mice. Science translational medicine 6:254ra127–254ra127.

64. Schindelin J, Arganda-Carreras I, Frise E, Kaynig V, Longair M, Pietzsch T, Preibisch S, Rueden C, Saalfeld S, Schmid B, Tinevez J-Y, White DJ, Hartenstein V, Eliceiri K, Tomancak P, Cardona A. 2012. Fiji: an open-source platform for biological-image analysis. Nature Methods 9:676–682.

65. Araujo P, Nguyen TT, Frøyland L, Wang J, Kang JX. 2008. Evaluation of a rapid method for the quantitative analysis of fatty acids in various matrices. J Chromatogr A 1212:106–13.

